# Redundant mechanisms driven independently by RUNX1 and GATA2 for hematopoietic development

**DOI:** 10.1101/2021.04.02.438148

**Authors:** Erica Bresciani, Blake Carrington, Kai Yu, Erika M. Kim, Tao Zhen, Victoria Sanchez Guzman, Elizabeth Broadbridge, Kevin Bishop, Martha Kirby, Ursula Harper, Stephen Wincovitch, Stefania Dell’Orso, Vittorio Sartorelli, Raman Sood, Paul Liu

## Abstract

RUNX1 is essential for the generation of hematopoietic stem cells (HSCs). *Runx1* null mouse embryos lack definitive hematopoiesis and die in mid-gestation. However, even though zebrafish embryos with a *runx1* W84X mutation have defects in early definitive hematopoiesis, some *runx1*^*W84X/W84X*^ embryos can develop to fertile adults with blood cells of multi-lineages, raising the possibility that HSCs can emerge without RUNX1. Here, using three new zebrafish *runx1*^*-/-*^ lines we uncovered the compensatory mechanism for *runx1*-independent hematopoiesis. We show that, in the absence of a functional *runx1*, a *cd41*-GFP^+^ population of hematopoietic precursors still emerge from the hemogenic endothelium and can colonize the hematopoietic tissues of the mutant embryos. Single-cell RNA sequencing of the *cd41*-GFP^+^ cells identified a set of *runx1*^*-/-*^-specific signature genes during hematopoiesis. Significantly, *gata2b*, which normally acts upstream of *runx1* for the generation of HSCs, was increased in the *cd41*-GFP^+^ cells in *runx1*^*- /-*^ embryos. Interestingly, genetic inactivation of both *gata2b* and its paralog, *gata2a*, did not affect hematopoiesis. However, knocking out *runx1* and any three of the four alleles of *gata2a* and *gata2b* abolished definitive hematopoiesis. *Gata2* expression was also upregulated in hematopoietic cells in *Runx1*^*-/-*^ mice, suggesting the compensatory mechanism is conserved. Our findings indicate that RUNX1 and GATA2 serve redundant roles for HSC production, acting as each other’s safeguard.

**Key points:**

- Existence of RUNX1-independent mechanisms for the generation of HSCs and the development of functional definitive hematopoietic cells
- GATA2 and RUNX1 functionally complement each other for their respective roles during hematopoiesis

## Introduction

The emergence and the maintenance of hematopoietic stem cells (HSCs) is regulated by several transcription factors including RUNX1 and GATA2^1-8^. *Gata2* is expressed in all functional HSCs and most of hematopoietic progenitor cells^9^. Mouse *Gata2*^*-/-*^ embryos are defective in definitive hematopoiesis and die at E10.5^6^. Moreover, *Gata2* is required for the generation of HSCs during the stage of endothelial to hematopoietic cell transition (EHT) in the aorta gonad mesonephric region (AGM)^8^. *GATA2* is also frequently involved in human leukemia, and germline mutations in *GATA2* are associated with an autosomal dominant disease with recurrent infections, myelodysplastic syndrome, and predisposition to leukemia^10^.

*Runx1*^*-/-*^ mice lack all definitive blood lineages and die in midgestation ^2^. *Runx1* is also required for HSC formation during embryo development^11^. Recent studies have shown that HSC precursors form but do not further develop into HSCs in *Runx1* deficient mice^12^. Our group previously generated a zebrafish line with a truncation mutation in *runx1* (*runx1*^*W84X/W84X*^)^13,14^. Similar to *Runx1*^*-/-*^ mice, *runx1*^*W84X/W84X*^ embryos failed to develop definitive hematopoiesis. Surprisingly, 20% of the *runx1*^*W84X/W84X*^ larvae could recover definitive hematopoiesis and survive to adult^14^. These findings suggested that HSCs could form without RUNX1 in the zebrafish. However, given the nature of the W84X mutation and the fact that *runx1*^*W84X*^ mRNA was still expressed in the *runx1*^*W84X/W84X*^ fish, we could not exclude the possibility of functional RUNX1 protein production through a stop-codon readthrough mechanism during translation.

We believe that elucidating the mechanism of this RUNX1-independent hematopoiesis has far-reaching implications in developing therapeutics for acute myeloid leukemia with germline or somatic mutations in RUNX1. Therefore, we generated three new *runx1*^*-/-*^ lines to uncover the mechanism of this RUNX1-independent production of HSCs. We detected *cd41*:GFP^+^ HSCs/HSPCs in the *runx1*^*-/-*^ embryos and characterized their expression profile by single-cell RNA sequencing (scRNA-seq). Interestingly, our analysis identified *gata2b* as one of the signature genes of *runx1-*independent HSC/HSPCs. We generated new *gata2b* and *gata2a* mutant lines and crossed them to the *runx1*^*-/-*^ lines to probe the interplay between RUNX1 and GATA2. Our findings suggest that RUNX1 and GATA2 act redundantly to support the formation and maintenance of HSCs and a functional hematopoietic system.

## Methods

### Zebrafish lines and maintenance

All zebrafish experiments were performed in compliance with NIH guidelines for animal handling and research. Zebrafish handling and breedings were performed as described previously^15^. The following zebrafish strains were used: wildtype (WT) zebrafish strain EK (Ekkwill) and TAB5, Tg(*gata1:*dsRed)^16^, *Tg(cd41*:GFP)^17^, *Tg(kdrl:mCherry)*^18^.

### *runx1, gata2a* and *gata2b* mutant generation and genotyping

Zebrafish mutants were generated with TALENS (*runx1* ^*del8*^ and *runx1* ^*del25*^) or CRISPR/Cas9 (*runx1* del(e3-8), *gata2a*, and *gata2b* mutants). The TALEN and sgRNA sequences, target exons and genotyping primer sequences for all mutations are described in Supplemental Table 1 and in the Supplemental Methods.

### Kidney RNA extraction and RNA sequencing

Kidneys were dissected from ∼2.5 months fish. See Supplemental Methods for detailed dissociation protocol. RNA-sequencing was performed on the HiSeq 2500 (Illumina) using paired-end library preparation (TruSeq RNA Library Prep Kit v2) at the NIH Intramural Sequencing Center. Low quality sequencing reads were removed with in-house script. RNA-Seq reads were aligned to the zebrafish genome reference (GRCz11) and transcript reference (GRCz11.99) using hisat2 (v2.2.1.0)^19^ on NIH HPC Biowulf cluster (http://hpc.nih.gov). We used htseq-count(v0.11.4)^20^ to generate gene expression estimates. DESeq2^21^ (version 3.4.2, http://www.r-project.org/) was used for differential gene expression analysis. Gene ontology enrichment analysis was performed using Metascape (http://metascape.org)^21^. Raw data has been deposited to GEO (GSE158098 and GSE169689).

### FACS-sorting and single cell capture and sequencing with 10X Genomics Chromium

Wildtype and *runx1*^*del8/del8*^ embryos were collected at 2.5 dpf, 6 dpf, 10 dpf, and 16 dpf and GFP+ cells were sorted on a FACS AriaIII (Becton Dickinson, Franklin Lakes, NJ). GFP^low^ cells were loaded on a Chromium Instrument (10x Genomics, Pleasanton, CA, USA) to generate single-cell GEMs. Single-cell RNA-seq libraries were prepared using Chromium Single Cell 3’ Library &Gel Bead Kit v3 (P/N1000075, 10x Genomics). The barcode sequencing libraries were sequenced on Illumina HiSeq2500/HiSeq3000.

### Data processing, clustering and trajectory analysis

Sequences from the Chromium platform were de-multiplexed and aligned using CellRanger ver. 2.0.2 from 10x Genomics using zebrafish reference genome (GRCz10) and transcript annotation reference (GRCz10.87). Clustering, filtering, variable gene selection and dimensionality reduction were performed using the Seurat ver.3.1.4^22,23^. Raw data matrices of two or more Seurat objects were merged to generate a new Seurat object with the resulting combined raw.data matrix. The cell identities of the clusters were re assigned based on the expression of well established hematopoietic markers that were identified by our unbiased analysis as cluster signature genes. The expression of signature markers that identify different cell identities is presented, for each time point, in heatmaps and feature maps (Fig. 3, 4 and Supplemental Fig. 2-4).

We applied STREAM (Version 1.0)^24^ software with single-cell gene expression matrix exported from Seurat to reconstruct the trajectories of different cell types. Different correlation metrics were used to calculate the ranks of cells, which reflect the cell position over the pseudotime axis. scRNA-seq data has been deposited in GEO (GSE158099). More details about data processing, clustering and trajectory analysis can be found in the Supplemental Methods.

### Mouse cell preparation and RNA-sequencing

All mice and procedures used in this study were approved by the National Human Genome Research Institute Animal Care and Use Committee. *Runx1* conditional knockout (*Runx1*^*f/f*^)^25^, *Mx1-Cre*^26^, and β-actin-Cre recombinase transgenic (Actb-Cre+)^27^ mice have been described previously. RNA-sequencing was performed on messenger RNA (mRNA) isolated from c-Kit^+^ bone marrow cells of mice^28^. Quantitative PCR (qPCR) was performed on mRNA extracted from whole Aorta-Gonad-Mesonephros (AGM) of E10.5 embryos. Details on the procedures and data analysis are provided in Supplemental Methods. Sequencing data has been deposited at Gene Expression Omnibus (GSE158100).

## Results

### Demonstration of RUNX1-independent hematopoiesis in three new *runx1* mutant lines

In order to investigate if RUNX1-independent emergence of HSCs exists, we generated three new *runx1*^-/-^ lines (Fig.1A and Supplemental Fig. 1A-D). Using transcription activator-like effector nucleases (TALENs) we obtained zebrafish lines with a 8 bp deletion and a 25 bp deletion (*runx1*^*del8*^ and *runx1*^*del25*^) within exon 4 of *runx1* (Fig. 1A, Supplemental Fig.1A and Supplemental Table 1). These mutations are predicted to cause frameshifts and premature terminations within the runt-homology domain (RHD), resulting in loss of function (Supplemental Fig. 1B and Supplemental Table 2). Since *runx1* mRNA was still detectable in *runx1*^*del8/del8*^ and *runx1*^*del25/del25*^ embryos (Supplemental Fig. 1C), we used two CRISPR-guide RNAs (Supplemental Table 1), targeting exon 3 (encoding RHD) and exon 8 (the last coding exon of *runx1*), respectively, to establish a mutant line carrying an exon 3-8 deletion (*runx1*^*del(e3-8)/del(e3-8)*^; Fig. 1A, Supplemental Fig. 1A, B, D and Supplemental Table 2), in which *runx1* full length transcript was not detectable (Supplemental Fig. 1C, D).

**Figure 1.**
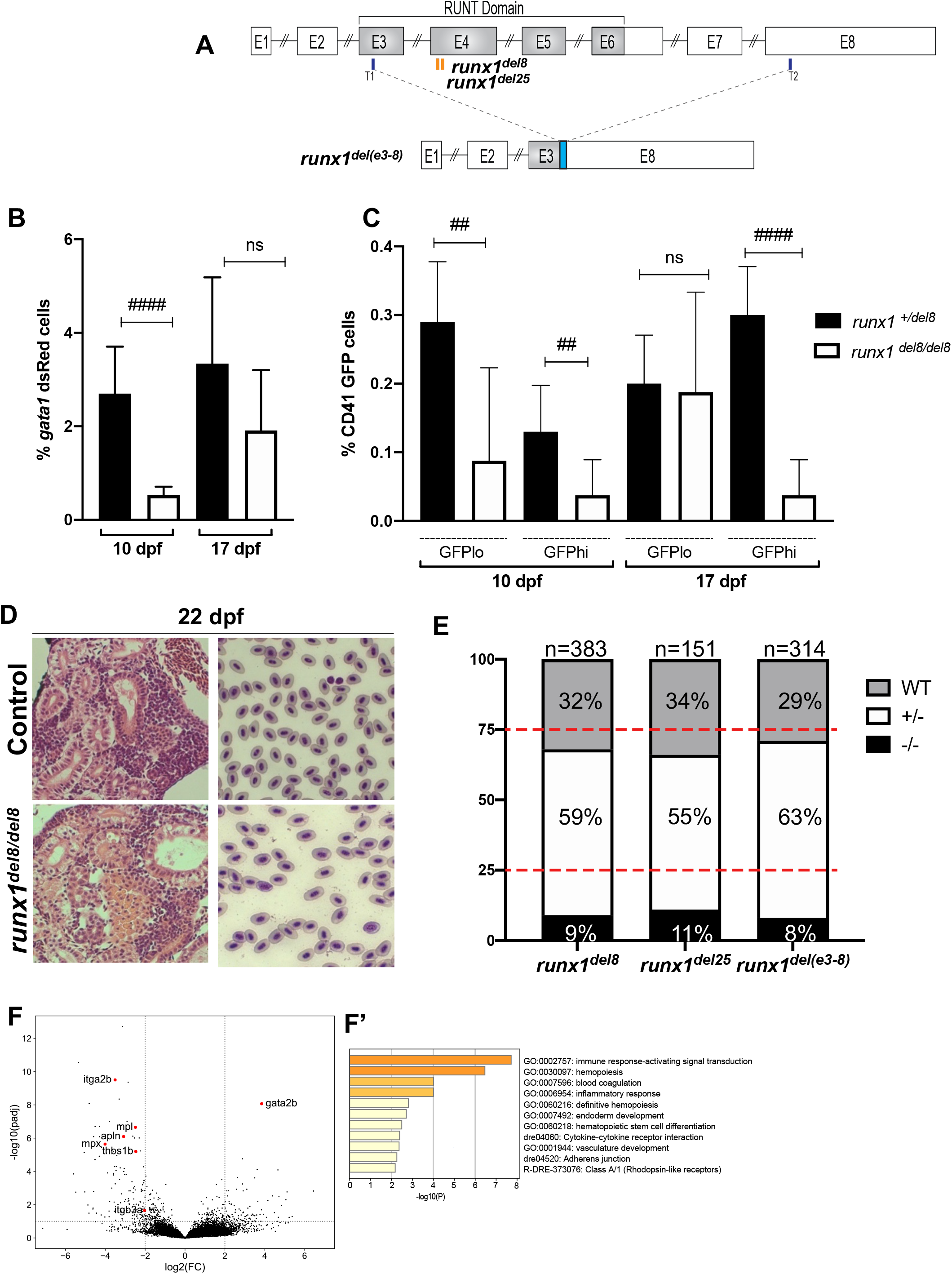
*runx1*^*-/-*^ mutants survive to adult and recover multi-lineage hematopoiesis. **A**. Schematic representation of the *runx1* gene showing the regions (orange bars) targeted with engineered TALENs that generated the *runx1*^*del8*^, runx1^del25^ mutant lines and the CRISPR targets, T1 and T2 (blue bars), used to generate the *runx1*^*del(e3-8)*^ mutant line as shown in the lower panel. The cyan box in the lower panel indicates a 66-bp insertion between the two cut sites. **B-C**. Percentage of *gata1:dsRed*^+^ (B) and *cd41*:GFP^low^ or *cd41*:GFP^high^ (C) in Tg(*gata1:dsRed; cd41:GFP*) *runx1*^*del8/del8*^ *runx1*^*+/del8*^ controls at 10 and 17 dpf analyzed by flow cytometry (##: p<0.01, ####: p<0.0001). **D**. Kidney histology and blood smear of a representative 22 dpf *runx1*^*del8/del8*^ appear similar to a *runx1*^*+/del8*^ control sibling. **E**. Stacked bar chart showing the percentage of adult *runx1*^*del/del8*^, *runx1*^*del2/del25*^, *runx1*^*del(e3-8)/ del(e3-8)*^ recovered from inbred heterozygous parents; red dashed lines indicate the expected Mendelian ratio. **F**. Volcano plot showing the differentially expressed genes in the kidneys between adult *runx1*^-/-^ and wildtype zebrafish. The red dots identify known hematopoietic markers for thrombocytes, myeloid cells and HSCs that are differentially expressed at FC>2 and padj <0.05. **F’**. Gene ontology enrichment analysis results for panel F.

Whole mount *in situ* hybridization showed lack of expression of all definitive hematopoietic markers (*c-myb, mpx, hbae1, rag1*) between 3-5 dpf in these three *runx1* mutant lines (Supplemental Fig.1E), indicating complete failure of definitive hematopoiesis at this stage. The larvae from all three *runx1* mutant lines were followed and found to undergo a bloodless phase between 10-16 dpf, during this time some of the *runx1* mutant larvae died. Circulating blood cells started to reappear in the remaining *runx1* mutant larvae between 16-20 dpf and by 22 dpf all the surviving *runx1* mutant larvae had blood circulation. The hematopoietic defects in these three mutant lines were qualitatively and quantitatively similar with each other. For practical reasons, we focused our subsequent studies on the *runx1*^*del8*^ mutant line.

Flow cytometric analysis showed significant reductions of *dsRed*^*+*^ erythrocytes and GFP^high^ thrombocytes in the Tg(*gata1:dsRed; cd41:GFP*) *runx1*^*del8/del8*^ larvae at 10 dpf (Fig. 1B and C). Interestingly, the GFP^low^ HSC/HSPC population was decreased in the mutants but still detectable (Fig. 1C). At 17 dpf, when circulating blood cells started to reappear in the *runx1*^*del8/del8*^ larvae, *dsRed+* erythrocytes and GFP^low^ HSC/HSPCs increased correspondingly but not the GFP^high^ thrombocytes (Fig 1 B, C). At 22 dpf, the kidneys in the *runx1*^*del8/del8*^ larvae were filled with blood progenitors, comparable to the controls and blood smear showed normal erythroid cells (Fig. 1D). Eventually, 8-11 % of the adult fish from *runx1*^+/-^ incrosses were *runx1*^-/-^ (Fig. 1E). *runx1*^-/-^ adults had normal size and morphology (Supplemental Fig.1F). Overall our data strongly suggests that in the absence of RUNX1 there is an independent mechanism for HSC formation.

### Gene expression changes in adult *runx1* null hematopoietic cells

To assess gene expression profile of hematopoietic cells in *runx1*^-/-^ adults we performed RNA-seq on dissected kidneys from both *runx1*^*del8*^ and *runx1*^*del25*^ (∼2.5 months old, n= 3 *runx1*^*del25*^, n=3 wild type siblings; n=2 *runx1*^*del8*^ and n=3 wild type siblings). Only 140 differentially expressed (DE) genes (107 downregulated and 33 upregulated, padj <0.05, FC>2) were detected between mutant and wildtype (Fig. 1F and Supplemental Table 3). Gene ontology (GO) enrichment analysis showed over-representation of genes related to immune response signaling/ inflammation and blood coagulation and hematopoiesis (Fig. 1F’). Indeed, key thrombocyte genes such as *itga2b* (*cd41*), *mpl, thbs1b, apln*, and *itgb3a* (*cd61*)^29^, were downregulated, indicating defects in thrombopoiesis in *runx1*^-/-^ fish (Fig. 1F). The myeloid markers *mpx* and *lyz* were also downregulated, suggesting an impairment of neutrophils. Interestingly, the HSC transcription factor *gata2b* was upregulated in *runx1*^*-/-*^ fish (Fig. 1F). Our results suggest that adult hematopoiesis in *runx1*^*-/-*^ fish is largely intact and multi-lineage.

### *cd41-*GFP^+^ HSC/HSPC are formed in *runx1*^-/-^ embryos

Our analysis of Tg(*gata1:dsRed; cd41:GFP*) *runx1*^*del8/del8*^ embryos showed the presence of cd41:GFP^low^ HSC/HSPCs at 10 dpf which were expanded during hematopoietic recovery. To identify where and when *runx1-*independent hematopoietic precursors might arise we generated *runx1*^*del8/del8*^; *Tg(cd41:GFP); Tg(kdrl:mCherry)* embryos. Interestingly, at ∼52 hpf, we could detect *cd41*-GFP^+^ mCherry^+^ cells in the AGM of *runx1*^*del8/del8*^ embryos, indicating that these cells were derived from hemogenic endothelium (Fig. 2A). Moreover, time-lapse confocal analysis showed that the *cd41-*GFP^+^ cells in the *runx1*^*del8/del8*^ embryos retained HSC behavior as they were released from the AGM into the circulation, similar to *cd41-*GFP^+^ HSCs in the wild type embryos (Supplemental Movies 1 and 2). At 3 dpf *cd41-*GFP^+^ cells had colonized the CHT of the *runx1*^*del8/del8*^ embryos (Fig. 2B), however, residual GFP^+^ cells could be found in the AGM (Fig. 2C) suggesting that the emergence/release of the HSPC in *runx1*^*-/-*^ was slower than normal. In addition, flow cytometric analyisis showed that the number of *cd41*-GFP^low^ cells at 2.5 dpf in the *runx1*^*del8/del8*^ was lower than normal (Supplemental Fig. 2A and B).

**Figure 2.**
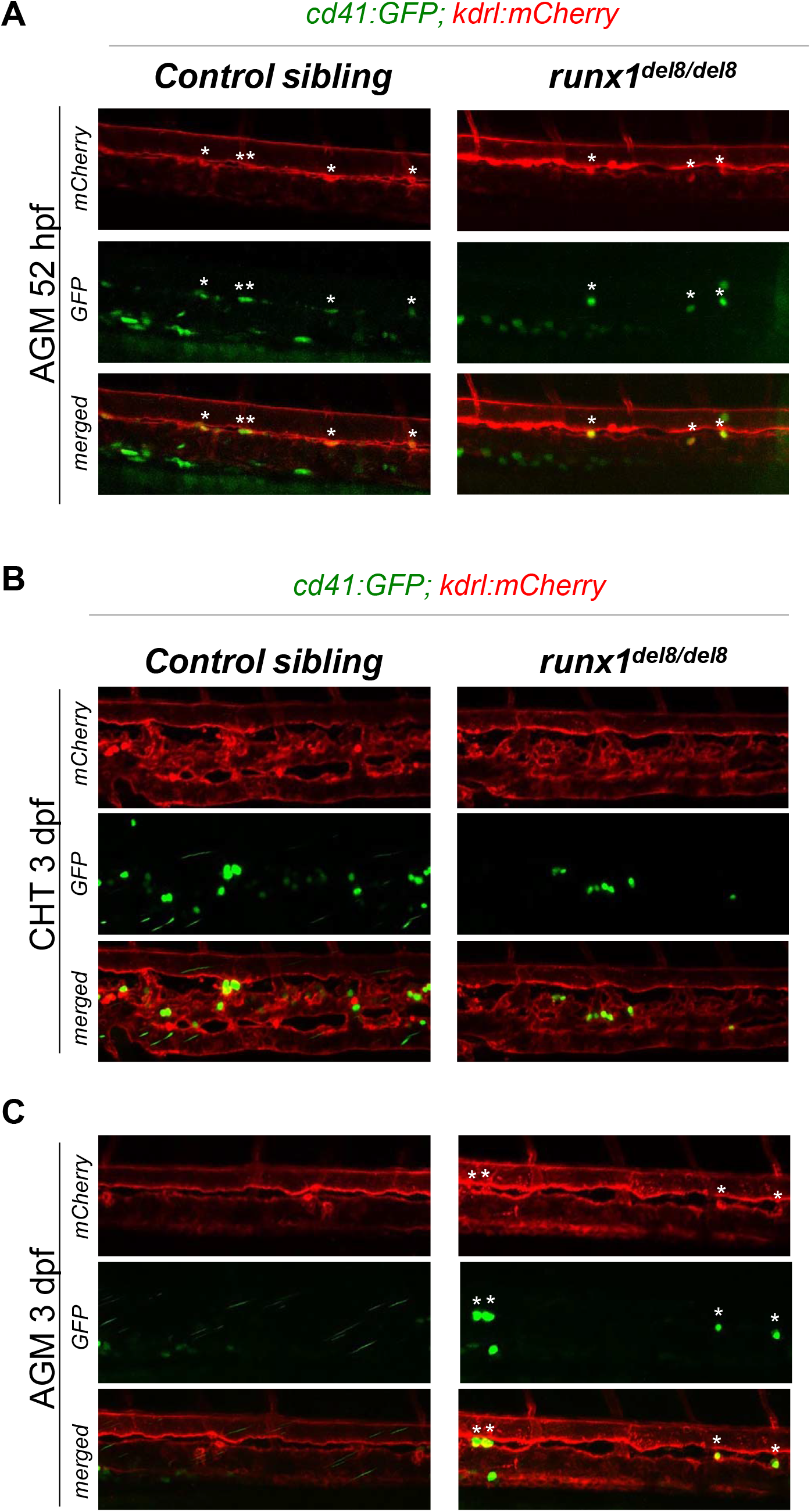
Development of *cd41*:GFP^+^ hematopoietic precursors in *runx1*^*del8/del8*^ AGM and CHT. **A**. Snapshots of the AGM region of *runx1*^*del8/del8*^ and control sibling *Tg(cd41:GFP); Tg(kdrl:mCherry)* at 52 hpf, obtained from time lapse recordings on a confocal microscope (Supplemental Movies 1 and 2). *cd41*:GFP^+^ mCherry^+^ double positive cells are indicated with white asterisks. **B-C**. Live confocal imaging of the CHT (B) and AGM (C) regions in the *Tg(cd41:GFP); Tg(kdrl:mCherry)* embryos at 3 dpf. *cd41*:GFP^+^ cells are found in the CHT (B). Residual *cd41*:GFP^+^ cells (white asterisks) are still detectable in the AGM of *runx1*^*del8/del8*^ (C).

The transcriptional profile of wildtype and *runx1*^-/-^ HSC/HSPCs was analyzed by scRNA-seq on *cd41*-GFP^low^ cells at 2.5 dpf (Supplemental Fig. 2C). Uniform Manifold Approximation and Projection (UMAP) of merged wildtype and *runx1*^*del8/del8*^ *cd41*-GFP^low^ cells defined 11 clusters (Fig. 3A and Supplemental Table 4). Cell identities were assigned to each cluster based on well established lineage markers identified as clusters signatures (Supplemetal Table 4). While the non-hematopoietic clusters were composed by mixed genotypes, the hematopoietic clusters were mainly composed by wildtype cells, indicating a reduction in the hematopoietic *cd41-* GFP^low^ cells in the *runx1*^-/-^ (Fig. 3A, B). The hematopoietic sub-populations expressed known markers for HSCs/HSPCs, multipotent progenitors, as well as erythroid and myeloid progenitors which likely represent erythro-myeloid progenitors (EMPs) generated at earlier stages of development and which are known to be *cd41:*GFP^*+30*^ (Fig. 3C, D and Supplemental Fig 2D).

Interestingly, both clusters 2 and 8 expressed HSC/HSPC signature genes such as *fli1a, tal1* (also known as *scl), c-myb*, and *lmo2* (Fig. 3A, C, D and Supplemental Fig. 2D). These two populations segregated based on their genotype with cluster 2 mainly composed of wildtype and cluster 8 mainly *runx1*^*del8/del8*^ (Fig. 3A, B). Only 41 DE genes were identified between clusters 2 and 8 (FC>2, padj<0.001; Supplemental Table 4), suggesting that the *runx1*^*del8/del8*^ HSC/HSPC population was not too different from the wildtype.

**Figure 3.**
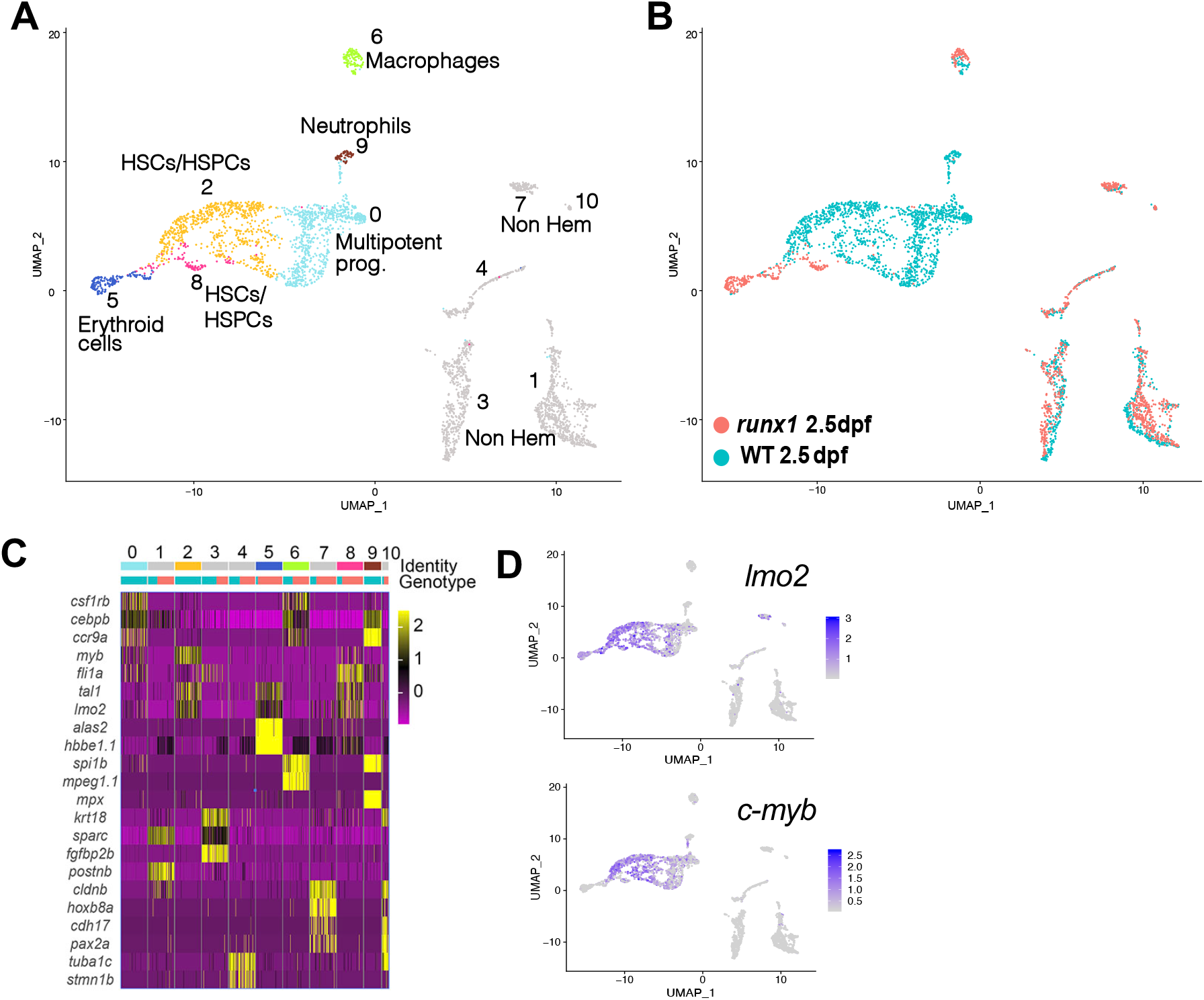
Single cell gene expression profile of wildtype and *runx1*^*del8/de*^ *cd41*:GFP^low^hematopoietic precursors at 2.5 dpf. **A**. UMAP of freshly FACS-isolated *cd41*:GFP^low^ cells from wildtype and *runx1*^*del8/del8*^ embryos at 2.5 dpf. Colored clusters represent hematopoietic cells, grey clusters are non-hematopoietic (based on expression profile). **B**. UMAP depicting the genotypes of the *cd41*:GFP^low^ cells. Blue dots, wildtype; Red dots, *runx1*^*del8/del8*^. **C**. Heat map depicting the expression of signature genes representative of different cell identities.The horizontal bars on the top correspond to the clusters identified in A (Identity) and show the distribution of the two genotypes (Genotype) across the clusters. 100 representative cells per clusters are shown. **D**. Feature plots depicting the expression level of the HSC markers *c-myb* and *lmo2* (purple is high, grey is low). See also Supplemental Fig.2 for additional scRNA-seq data from 2.5 dpf embryos.

### *runx1*^*-/-*^ HSCs/HSPCs at larval stages had distinct gene expression profile

To uncover the compensatory mechanism that drive the hematopoietic recovery in *runx1*^-/-^ we characterized the expression profile of *cd41*-GFP^low^ cells during larval development by scRNA-seq (Supplemental Fig. 3A, B). UMAPs of merged wildtype and *runx1*^-/-^ at 6 dpf identified multiple hematopoietic subpopulations, highlighting the level of heterogeneity of *cd41-*GFP^low^ cells at this stage (Fig. 4A, Supplemental Table 5). HSCs/HPSCs (clusters 1, 3, 5) expressing markers such as *tal1, lmo2* and *c-myb* (Fig. 4C and Supplemental Fig. 3C) again segregated based on genotype (clusters 1 wildtype; cluster 3 mutant; Fig. 4A-C). Cluster 5, again predominantly wild type, was composed by HSCs undergoing cell division as shown by the specific expression of cell cycle markers *ccna2* and *ccnb1* (Fig. 4C). Notably, *meis1b, fli1a* and *gata2b* were among the signature markers for the *runx1*^*del8/del8*^ HSC/HSPC (Fig. 4C and Supplemental Fig. 3C).

**Figure 4.**
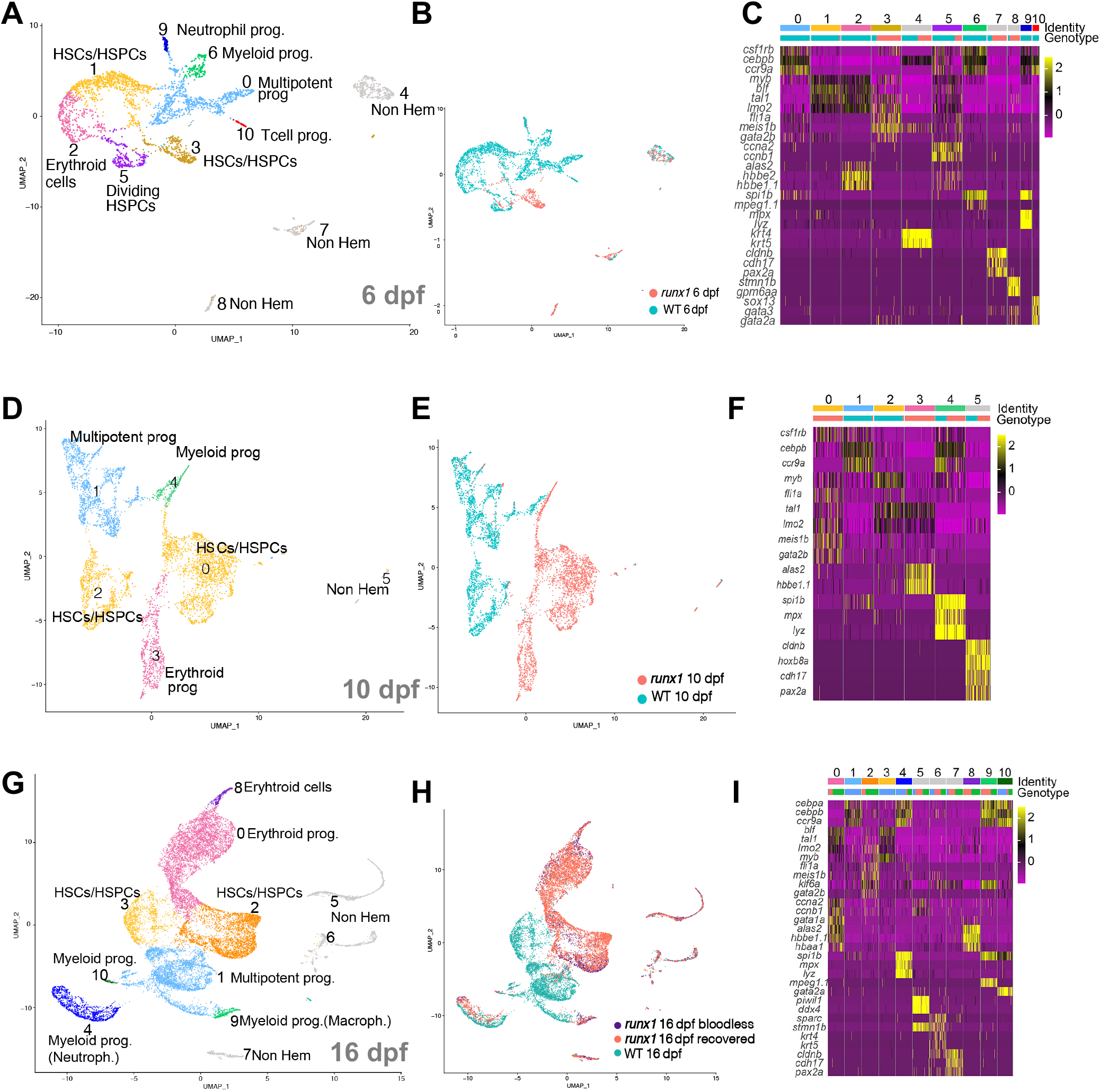
Single cell analysis of wildtype and *runx1*^*del8/del8*^ *cd41*:GFP^low^ at 6, 10 and 16 dpf show only few overlapping populations. **A, D, G**. UMAP of freshly FACS-isolated *cd41*:GFP^low^ cells from wildtype and *runx1*^*del8/del8*^ embryos at 6 dpf (A), 10 dpf (D) and 16 dpf (G). Colored clusters represent hematopoietic cells, grey clusters are non-hematopoietic (based on expression profile). **B, E, H**. UMAP depicting the genotypes of the *cd41*:GFP^low^ cells at 6 dpf (B), 10 dpf (E) and 16 dpf (H). Blue dots, wildtype; Red dots, *runx1*^*del8/del8*^ (B, E). At 16 dpf Blue dots represent wildtype; Purple dots, *runx1*^*del8/del8*^ with circulating blood cells (recovered), Red dots, bloodless *runx1*^*del8/del8*^. **C, F, I**. Heat maps depicting the expression of signature genes representative of different cell identities in each of the clusters identified respectively in A, D, G. The horizontal bars on the top correspond to the clusters identified the correspondent UMAP (Identity) and show the distribution of the two genotypes (Genotype) across the clusters. 100 representative cells per clusters are shown. See also Supplemental Fig. 3 and 4 for additional scRNA-seq data.

At 10 dpf, wildtype and mutant *cd41-*GFP^low^ cells again clustered separately (Fig. 4E). For the first time, the *runx1*^*del8/del8*^ cells showed a higher degree of heterogeneity with *runx1*^*del8/del8*^ specific HSC/HSPC and erythroid progenitors clusters, and one myeloid progenitors population shared with wildtype (Fig. 4D-F and Supplemental Fig. 3D). At this stage, *meis1b, fli1a* and *gata2b* remained specifically expressed in the *runx1*^*del8/del8*^ HSC/HSPCs (Fig. 4F, Supplemental Fig. 3D and Supplemental Table 5).

At 16 dpf, the *runx1*^-/-^ larvae were separated based on their phenotype: “bloodless” (without circulating blood cells) and “recovered” (regained some or nearly normal levels of circulating blood cells). This large dataset of ∼11k *runx1*^*del8/del8*^ cells presented five hematopoietic populations and showed complete overlap between “bloodless” and “recovered” (Supplemental Fig. 4A-C and Supplemental Table 5), suggesting these larvae had the same hematopoietic progression despite of the phenotypic difference.

Analysis of a merged dataset containing wildtype and *runx1*^*del8/del8*^ (bloodless and recovered) *cd41*-GFP^low^ cells at 16 dpf showed again that most cells clustered based on their genotypes (Fig. 4G-I, Supplemental Table 5). Importantly, the wildtype and *runx1*^*del8/del8*^ HSC/HSPC clusters were distinct from each other. There were two *runx1*^*del8/del8*^ specific erythroid clusters (clusters 0 and 8) expressing different levels of *gata1a, urod*, hbae1, *hbaa1*, and *alas2* (Fig. 4G-I and Supplemental Fig. 4D). On the other hand, multipotent progenitor cluster (cluster 1) was mostly composed of wildtype cells. Only the myeloid clusters 4 and 10 were shared by wildtype and *runx1*^*del8/del8*^ cells (Fig. 4G-I and Supplemental Fig. 4D). Overall our data suggest that the HSC population in the *runx1*^*del8/del8*^ larvae, though different from the wildtype, could give rise to erythroid and myeloid progenitors but a multi-progenitor population was not established at this stage.

### Comparing the hematopoietic differentiation trajectories in *runx1*^*del8/del8*^ and wildtype larvae

With the aim to reconstruct the progression of the hematopoietic recovery in the *runx1*^*del8/del8*^larvae, we merged scRNA-seq datasets from 6, 10, and 16 dpf based on genotype (Supplemental Fig. 5A, B, D, E and Supplemental Table 6) and performed trajectory analysis (STREAM^24^).

The unbiased trajectory reconstruction of wildtype *cd41* GFP^low^ cells (11,683 cells) identified 8 different branches (Fig. 5A, and Supplemental Fig. 5C). The HSC/HSPCs were split in 3 arms (arms S2-S0, S2-S4, S2-S5, Fig. 5A, B and Supplemental Fig. 5C). Arm S2-S0 was composed largely by HSCs/HSPCs at 10 and 16 dpf; S2-S4 was populated by HSCs/HSPCs from 10 dpf differentiating into erythroid progenitors expressing *gata1, alas2* and embryonic globins (*e*.*g. hbbe1;* Fig. 5A, B and Supplemental Fig. 5C, G). Arm S2-S5 cells were exclusive of 16 dpf (Fig. 5A, B) and expressed adult globin *hbaa1*, suggesting these cells were responsible for the switch from embryonic to adult erythropoiesis^31^ (Fig. 5A, B and Supplemental Fig. 5C, G). Cells in the S0-S1 branch were derived from *cd41*-GFP^low^ at 6 dpf and expressed embryonic erythroid markers (Fig. 5A, B and Supplemental Fig. 5C, G). The remaining branch of HSPCs (S0-S3) differentiate toward myeloid progenitors with macrophage signatures (S3-S7), neutrophils (S3-S8) and B/T progenitor cells (S3-S6) (Fig. 5A, B and Supplemental Fig. 5C, G).

**Figure 5.**
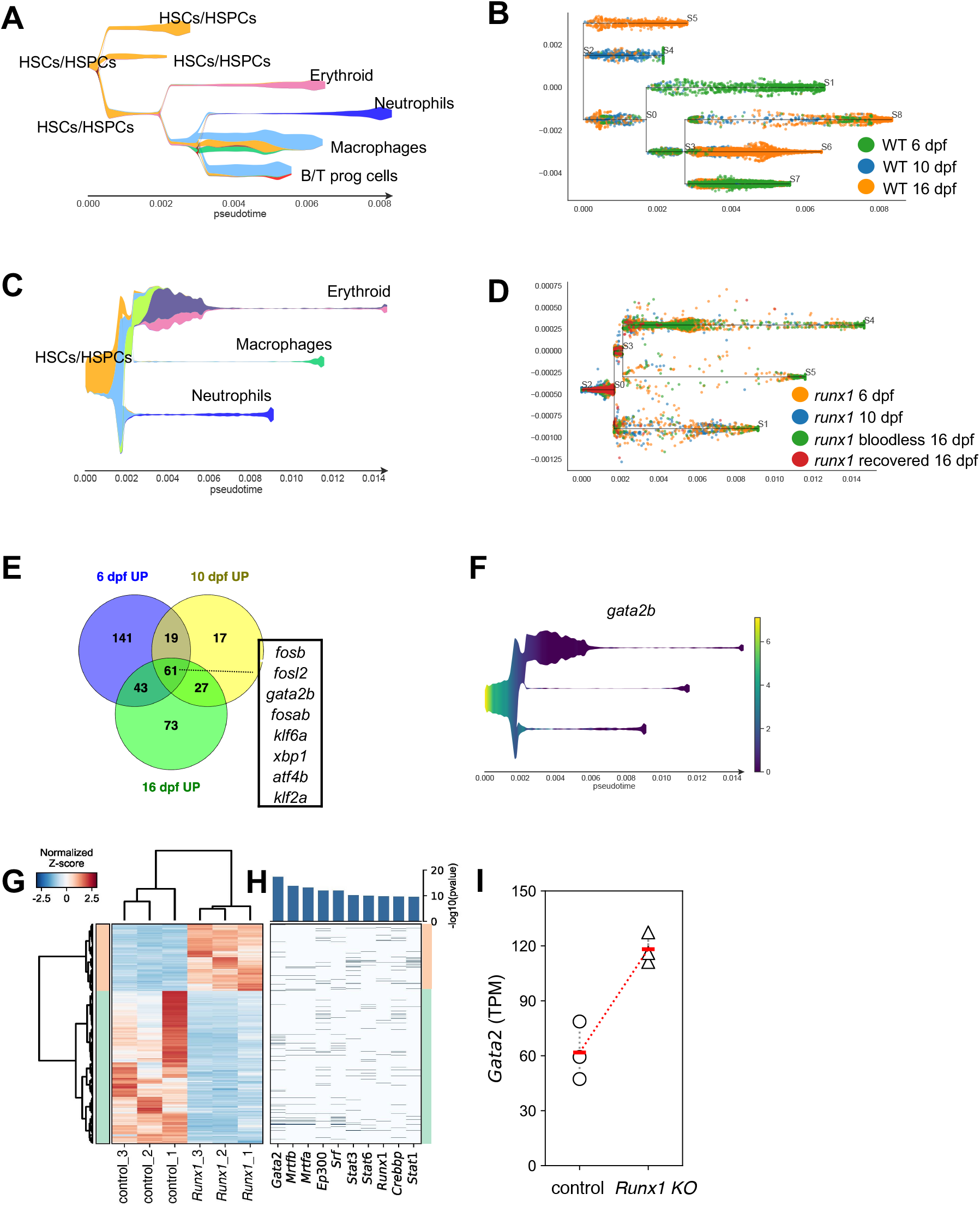
*Gata2* is up-regulated in the HSPCs of *Runx1* zebrafish and mice knockouts. **A**. Stream plot representing the pseudotime trajectory projection of wildtype *cd41*:GFP^low^ cells at 6, 10, 16 dpf and their different identities. **B**. Subway map depicting the distribution of wildtype *cd41*:GFP^low^ cells from different time points to the different braches. **C**. Stream plot representing the pseudotime trajectory projection of *runx1*^*del8/del8*^ *cd41*:GFP^low^ cells at 6, 10, 16 dpf and their different identities. **D**. Subway map illustrating the contribution of *runx1*^*del8/del8*^ *cd41*:GFP^low^ at different time points to the different braches. **E**. Venn diagram representing the number of upregulated genes in *runx1*^*del8/del8*^ HSC/HSPCs vs wildtype at 6, 10,16 dpf (padj<0.05, FC>1.5). 61 genes were commonly upregulated, of which 8 were transcription factors (listed). **F**. Stream plot depicting the expression of *gata2b* in the *runx1*^*del8/del8*^ larval cd41-GFP^low^ pseudo-time development (from panels C and D). See also Supplemental Figure 5 for additional analyses of hematopoietic differentiation trajectories in *runx1*^*del8/del8*^ and wildtype larvae at 6, 10, and 16 dpf. **G**. Unsupervised hierarchical clustering of wildtype and *Runx1*^*-/-*^ RNAseq samples and heat map depicting the differentially expressed genes between wildtype and *Runx1*^*-/-*^ c-Kit^+^ HSPCs (940 downregulated and 414 upregulated in the *Runx1*^*-/-*^; padj<0.05, FC>2). **H**. Top transcription factors enriched for regulating the differentially expressed genes. The X-axis of the heatmap lists the transcription factors and the Y-axis are the DE genes sorted in the same order of the Y-axis in B. Barplot on the top showed the enrichment p-values for each transcription factor. **I**. *Gata2* expression in c-kit+ bone marrow cells in control and *Runx1*^*-/-*^ mice, measured by RNA-seq. Round and triangle points mark TPM values of each mouse in control and *Runx1*^*-/-*^ mice, respectively. Red ticks mark the average TPM value. See also Supplemental Figure 6 for additional data on *Gata2* expression in *Runx1* conditional knockout mice.

The trajectory of *runx1*^*del8/*^ cells (16,660 cells) showed a lower number of bifurcations compared to the wildtype (Fig. 5C, D and Supplemental Fig. 5F). The *runx1*^*del8/del8*^ cells predominantly populated the most undifferentiated branches: HSCs/HSPCs (S2-S0), HSPCs/Multipotent progenitors (S2-S0, S0-S3, S0-S1) and HSPCs, which were undergoing active translation based on GO term analysis (“translating HSPCs”, S0-S3; Fig. 5C, D, Supplemental Fig. 5F, H and Supplemental Table 6). HSPCs/Multipotent progenitors and translating HSPCs also populated the erythroid trajectory branch (S3-S4) which includes erythroid progenitors with both embryonic and adult globin signatures (*hbbe1, hbbe2* and *hbaa1* respectively), and the myeloid branches with neutrophils (S0-S1) and macrophages signatures (S0-S5; Fig. 5C, D, Supplemental Fig. 5F, H). Overall, the lower level of complexity and the preponderant distribution of the *runx1 cd41*-GFP^low^ cells in the more undifferentiated branches were consistent with delayed hematopoiesis in the *runx1*^*del8/del8*^ larvae.

### G*ata2* is up-regulated in the HSPCs in the *Runx1* knockout zebrafish and mice

In order to gain insight into the compensatory mechanism taking place in *runx1*^*-/-*^ HSCs/HSPCs we considered the upregulated genes in the *runx1*^*del8/del8*^ HSC/HSPCs at 6, 10 and 16 dpf (Fig. 5E and Supplemental table 5). We reasoned that another transcription factor was likely responsible for compensation of *runx1*. Among 61 genes consistently upregulated in *runx1*^*del8/del8*^ HSC/HSPCs (padj<0.05, FC>1.5) only 8 were transcription factors (Fig. 5E). Significantly, in the absence of a functional runx1, *gata2b* was the only transcription factor that remained persistently upregulated in the HSC population at larval stages and also in the adult kidney (Fig. 5E, F and 1F, F’).

We also examined the expression levels of *Gata2* in the *Runx1* knockout mice to determine if the interplay between *runx1* and *gata2* is evolutionarily conserved. *Gata2* expression level in the AGM of *Runx1* conditional knockout mouse embryos (*Runx1*^*f/f*^, β*-actin-Cre)* at E10.5 was unchanged (Supplemental Fig. 6A-C). We then examined the gene expression profiles in c-Kit^+^ bone marrow HSPCs from adult *Runx1* conditional knockout mice (*Runx1*^*-/-*^, *Mx1-Cre*)(Supplemental Fig. 6D)^25,32^. We identified a total of 1354 differentially expressed genes (padj<0.05, FC>2) between *Runx1*^*-/-*^ and control c-Kit^+^ HSPCs (Fig. 5G, Supplemental Fig. 6E and Supplemental Table 7), and Ingenuity Pathway identified GATA2 as the top upstream regulator responsible for the gene expression changes (Fig. 5H and Supplemental Fig. 6F). Moreover, *Gata2* expression was increased in the *Runx1*^*-/-*^ c-Kit^+^ HSPCs with a highly significant padj (9.41E-06, Fig. 5I). These findings suggest that the interplay between *Runx1* and *Gata2* is evolutionarily conserved.

### *gata2* is required for *runx1-*independent hematopoiesis

To determine if *gata2b* could compensate in the absence of *runx1* we generated two *gata2b* mutant lines (Supplemental Table 1), a 5 bp insertion and a 7 bp deletion located upstream and within the first zinc-finger domain of GATA2, respectively (Fig. 6A, Supplemental Fig. 7A, B and Supplemental Table 2). Surprisingly both *gata2b* mutants reached adulthood according to Mendelian ratio with no hematopoietic phenotype (Supplemental Fig. 7C). However, when *runx1*^*del8/del8*^ and *gata2b*^*ins5/ins5*^ fish were bred together, *runx1*^*del8/del8*^; *gata2b*^*ins5/ins5*^ adults were recovered at lower than expected number, suggesting that *gata2b* is important for the survival of the *runx1*^-/-^ fish (Fig. 6B and Supplemental Fig. 7D). On the other hand, injection of *runx1*^*del8/del8*^; *Tg(cd41:GFP)* embryos with a published *gata2b* splicing morpholino, *gata2b*-MO^33^, led to loss of *cd41*-GFP+ cells in AGM and CHT (n= 26/29, Supplemental Fig. 7E), and significant decrease of survival at 1 month after injection (Supplemental Fig. 7F).

**Figure 6.**
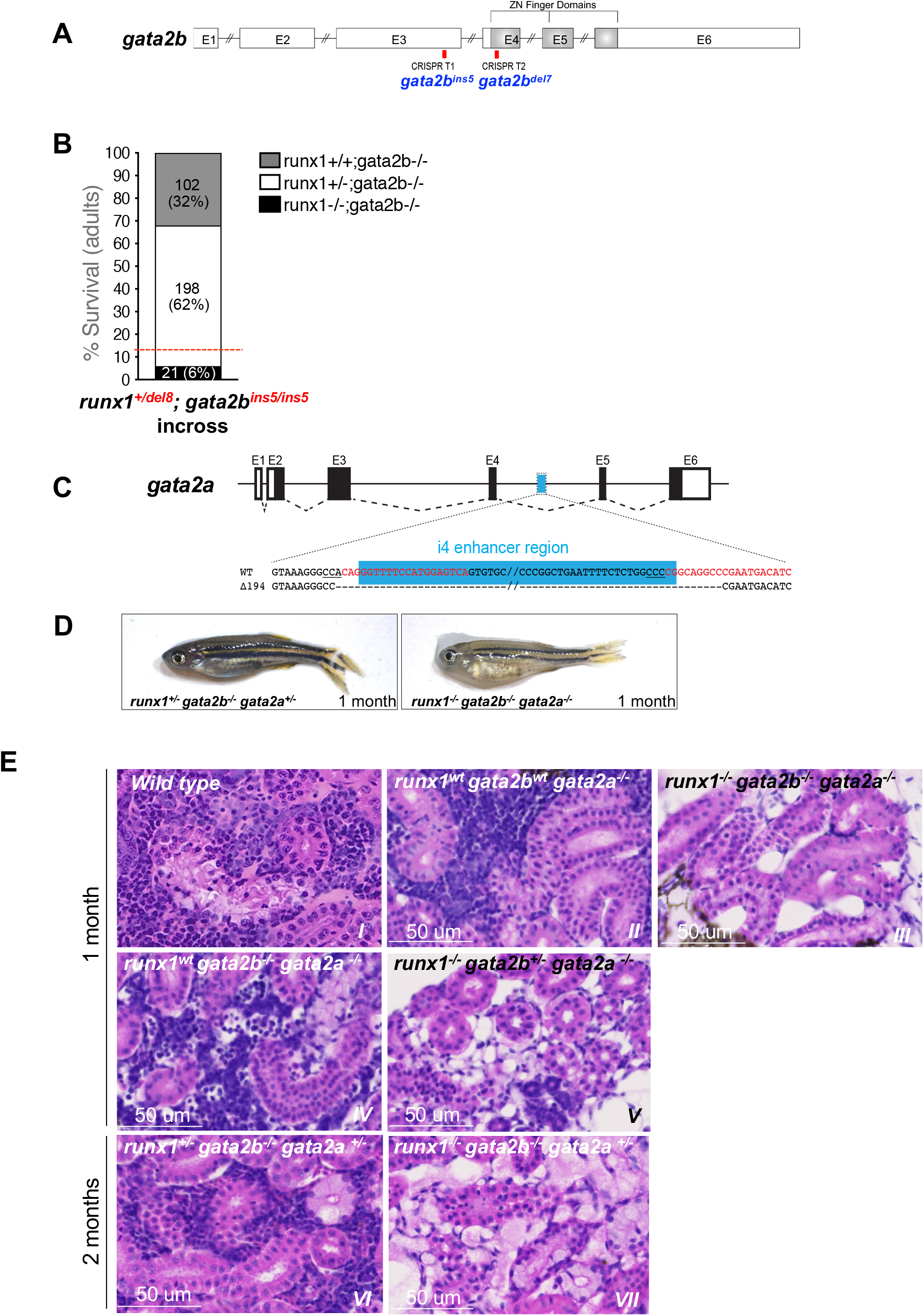
*gata2b* and *gata2a* are required for the development of definitive hematopoiesis in *runx1* mutants. **A**. Schematic representation of the *gata2b* gene showing the CRISPR targets T1 and T2 (red bars) used to generate the *gata2b*^*ins5*^ and the *gata2b*^*del7*^ mutant lines. **B**. Survival of adult *runx1*^*del8*^*/gata2b*^*ins5*^ double mutants obtained from the incross of *runx1*^*+/del8*^; *gata2b*^*ins5/ins5*^ fish. Red dashed lines indicate the expected ratio of *runx1*^*-/-*^ recovery based on our previous experimental data (Fig. 1E). Each bar segment shows the percentage and number of fish recovered for each genotype. **C**. Schematic representation of *gata2a* genomic structure. The cyan box represents the 150 bp i4 enhancer that was removed using two CRISPR guides (Red letters). **D**. Photos of representative 1 month-old *runx1*^*+/-*^ *gata2b*^*-/-*^ *gata2a*^*+/-*^ and *runx1*^*-/-*^ *gata2b*^*-/-*^ *gata2a*^*-/-*^ fish, with the latter more pale and smaller than the former. **E**. Histological analysis of kidney marrow of triple mutants *runx1*^*del8*^; *gata2b*^*ins5*^; *gata2a*^*i4del1*^ at 1 or 2 months. Kidneys from runx1 wildtype and *runx1*^*+/-*^ fish (panels *II, IV, VI*) are similar to the kidney from the wildtype control (panel I) with marrows populated by blood progenitors. Kidneys from *runx1*^*del8/del8*^ fish with *gata2b*^*-/-*^ *gata2a* ^*-/-*^ (*III*), *gata2b*^*+/-*^ *gata2a* ^*-/-*^ (*V*), or *gata2b*^*+/-*^ *gata2a*^*+/-*^ (*VII*) completely lack blood progenitors. See also Supplemental Figure 7 for additional characterizations of the zebrafish *gata2b* and *gata2a* mutants.

The normal survival and the absence of obvious hematopoietic phenotypes in both *gata2b* mutants raised the possibility of compensation by the paralogous gene, *gata2a*, which is also expressed in the aortic endothelium^34^. Using a published CRISPR pair^34^, we deleted the i4 enhancer of *gata2a* (*gata2a*^*i4del194*^) in *runx1; gata2b* fish, to reduce *gata2a* expression specifically in the hemogenic endothelium (Fig. 6C and Supplemental Fig. 7G). We assessed the hematopoietic phenotype in the triple mutant line (*runx1*^*del8*^, *gata2b*^*ins5*^, and *gata2a*^*i4del194*^). At 15 dpf, the bloodless phenotype was observed only in *runx1*^*del8/del8*^ larvae, irrespective of *gata2b* and *gata2a* mutation status (Supplemental Fig. 7H). However, at 1-2 month of age, the only fish that remained bloodless (and smaller than their clutchmates) were *runx1* ^*del8/del8*^ with at least 3 *gata2* mutant alleles (*gata2b*^*+/-*^*/gata2a*^-/-^, *gata2b*^*-/-*^*/gata2a*^*+/-*^, or *gata2b*^*-/-*^*/gata2a*^*-/-*^)(Fig. 6D). Moreover, histological analysis showed that all *runx1*^*-/-*^ fish with at least 3 *gata2* mutant alleles had no hematopoietic progenitors in their kidneys (Fig. 6E). These findings clearly indicated that *gata2a* and *gata2b* were required for the hematopoietic recovery in the *runx1* nulls. Interestingly, hematopoiesis appeared normal in the kidneys of *runx1*^*wt*^; *gata2b*^*-/-*^ ; *gata2a*^*-/-*^ fish. Even though we could not exclude the presence of lineage specific defects in the absence of GATA2, our data suggests that definitive hematopoiesis could proceed without *gata2* (Fig. 6E).

## Discussion

The three new zebrafish *runx1*^*-/-*^ lines and the previously reported *runx1*^*W84X/W84X*^ failed to initiate definitive hematopoiesis during early embryonic development, but between 30-40% of the *runx1*^*-/-*^ larvae could regain hematopoiesis and survive to fertile adults. These findings definitely proved the presence of a RUNX1-independent mechanism for HSC production and definitive hematopoiesis. Interestingly, the expression profile of *runx1*^*-/-*^ adult kidney cells was relatively normal except for the thrombocyte lineage which is dependent on RUNX1^14,35,36^.

We observed *cd41*:GFP^low^ cells in the AGM of *runx1*^*-/-*^ embryos at 2.5 dpf, which were derived from hemogenic endothelium and possessed HSC-like gene expression profile. *cd41*:GFP^low^ HSC/HSPCs were later expanded in the kidney of *runx1*^*-/-*^ larvae during the visible recovery of circulating blood cells. Therefore, we hypothesize that these *cd41*:GFP^low^ HSC/HSPCs are responsible for hematopoiesis recovery in *runx1*^*-/-*^ larvae.

We characterzed the *cd41*:GFP^low^ HSC/HSPCs in the *runx1*^*-/-*^ larvae by scRNA-seq. At all stages these cells differed significantly from wildtype cells in levels of heterogeneity and expression profile, with identification of signature genes unique to *runx1*^*-/-*^ *cd41*:GFP^low^ HSC/HSPCs. In fact, the cd41:GFP^low^ HSC/HSPCs in the *runx1*^*-/-*^ larvae clustered separately from the wild type HSC/HSPCs at all stages, suggesting that they have a unique aberrant phenotype, rather than having a developmental delay or block. Trajectory analysis across three larval stages demonstrated that *runx1*^*-/-*^ *cd41*:GFP^low^ cells were skewed towards earlier stages of development. At 10 and 16 dpf, contributions to the development of the erythroid and myeloid branches could be observed in the *runx1*^*del8/del8*^, probably representing the hematopoietic expansion during the recovery period. *runx1*^*del8/del8*^ larvae appeared to have a large number of *cd41*:GFP^low^ erythroid progenitors, even though cd41:GFP is not known to be expressed in erythroid cells^17,29,37^. We speculate that these erythroid progenitors were “recently” generated from the GFP^low^ HSC/HSPCs and therefore still retained GFP expression.

To uncover the mechanism for RUNX1-independent hematopoiesis we focused our attention on transcription factors differentially upregulated in the *runx1*^*-/-*^ HSC/HSPCs. *gata2b* was the only transcription factor upregulated at all larval stages and also in the kidney of *runx1*^*-/-*^ adults. Importantly, *gata2a* was not differentially expressed in the *runx1*^*-/-*^ at any time point. GATA2 is required for HSC development and is known to act upstream of RUNX1 in the hemogenic endothelium^8,38,39^. Zebrafish contains two *GATA2* paralogs, *gata2a*, which is expressed in the whole vasculature, included hemogenic endothelium^34^ and *gata2b*, which is specifically expressed in hemogenic endothelium^33^. A previous publication showed that, at the onset of definitive hematopoiesis, *gata2a*^Δ*i4/*Δ*i4*^ mutants present an initial delay in the induction of hemogenic endothelium but then can progress with a normal hematopoietic development^34^. *gata2b* mutants, on the other hand, present a reduction in HSC at the onset of definitive hematopoiesis and have defect in neutrphil and monocytic differentiation lineages as adults^40^. Surprisingly, there were no obvious hematopoietic defects in the adult *gata2b*^*-/-*^ ; *gata2a*^*-/-*^ fish, suggesting GATA2 is not required for hematopoiesis^34^. However, recovery of definitive hematopoiesis in *runx1*^*-/-*^ fish was completely abolished in the absence of at least 3 of the 4 *gata2a/2b* alleles. Our results show that GATA2 is responsible for RUNX1-independent hematopoiesis. Moreover, the results suggest that GATA2 and RUNX1 complement each other for their respective roles during hematopoiesis. Even though there have been reports that they work synergistically^41^, our findings suggest that they can work independently, providing redundant mechanisms for HSC production.

It has been reported that CD41+ pre-HSCs can be found in the ventral dorsal aorta of the *Runx1*^*ko*^ mice around E10.5^12^. Based on our analyses, *Gata2* did not appear to be upregulated in the AGM of the *Runx1*^*ko*^ mice. However, *Gata2* is upregulated and identified as the top upstream regulator of the differentially expressed genes in the c-Kit^+^ HSPCs in adult *Runx1*^*-/-*^ mice. Therefore, it is possible that a RUNX1-independent hematopoiesis initiates in both zebrafish and mouse, but this process is successful only in zebrafish because of its larvae can survive in the absence of blood circulation. Because of this mechanism, several other bloodless mutants in the zebrafish could be partially or completely rescued through compensating mechanisms^43-46^.

Moreover, RUNX1 and GATA2 may interact with each other for leukemia development. As transcription factors, RUNX1 and GATA2 can interact with each other directly and form DNA-binding complexes^41^. RUNX1 and GATA2 binding sites are found on a number of hematopoietic genes, and these two transcription factors have bee shown to cooperate with other transcrption factors to regulate genes considered critical for the balance between quiescence and proliferation of HSCs^41^, as well as genes involved in leukemia^47^. We recently found that *GATA2* is frequently deleted in human CBF leukemias, those that are associated with *RUNX1* and *CBFB* fusion genes^48^. In addition, we demonstrated that, indeed, *Gata2* contributes to leukemogenesis associated with *CBFB-MYH11* in a mouse model^49^. Interestingly, we have also demonstrated that *Runx1* is required for leukemogenesis associated with *CBFB-MYH11* in mouse models^50^, raising the possibility that *Gata2* and *Runx1* function synergistically during leukemogenesis.

Overall, our data suggest that GATA2 serves as a safeguard for the formation of definitive HSCs/HSPCs in case RUNX1 is inactivated and that this mechanism is evolutionary conserved. It may not be surprising that GATA2 serves this role since both GATA2 and RUNX1 are key transcription factors for hematopoiesis. The interaction between RUNX1 and GATA2 at common target genes has previously been reported, and interestingly *Gata2*^*+/-*^ *Runx1*^*+/-*^ mice are embryonic lethal with hematopoietic defects^41^. However, it is surprising that GATA2 can bypass RUNX1 to establish functional hematopoiesis. Likewise, RUNX1 seems to be able to sustain hematopoiesis in the absence of GATA2. Only when losing both RUNX1 and at least 75% of GATA2 was hematopoiesis abolished, suggesting remarkable redundancy between GATA2 and RUNX1 to ensure the initiation and maintenance of HSC that can support hematopoiesis, a vital system for the survival of vertebrate animals.

## Supporting information

Supplemental Movie 2 Mut

## Acknowledgements

We thank the NIH Intramural Sequencing Center (NISC) for RNA-seq of the kidney samples, and Brant Weinstein for the *Tg(kdrl:mCherry)* fish line. We also thank all the Liu lab members, Alberto Rissone and Aniket Gore for helpful discussions and advice. We thank Faiza Naz for her support with single cell cDNA synthesis. The work described in this paper was supported by the Intramural Research Programs at the National Human Genome Research Institute, NIH and the National Institute of Arthritis and Musculoskeletal and Skin Diseases, NIH. This work utilized the computational resources of the NIH HPC Biowulf cluster. (http://hpc.nih.gov)

## Authorship

### Contribution

E.B., B.C, K. B., V.S.G. and E.B. designed and performed the experiments and analyzed the data; E.B, R.S., and P.P.L. designed and organized the experiments and analyzed the data. M.K. assisted with fluocytometry, U.H. assisted with genotyping, S.W. assisted with microscopy. S.D.O. and V.S. assisted with capture and sequencing of 10X Genomics and provided technical support. T.Z. designed and performed mouse experiments. E.B, E.M.K. and K.Y. performed bioinformatic analysis. E.B. and P.P.L. wrote the manuscript.

