## Supplemental Methods for "Redundant mechanisms driven independently by RUNX1 and GATA2 for hematopoietic development"

***runx1*, *gata2a* and *gata2b* mutant generation and genotyping**

TALENs targeting exon 4 of *runx1* were assembled using the Golden Gate TALEN system^34,35^. Synthesized TALEN RNA (75pg left arm and 75pg right arm) was injected in zebrafish embryos and founder fish and F1 heterozygous adults were screened as previously described by fluorescent PCR^36^. CRISPRs for *runx1* del(e3-8), *gata2a* and *gata2b* exon 3 were synthesized using the clone free method as previously described^37^. Zebrafish embryos were injected with 50pg of sgRNA(s) mixed with 300pg of Cas9 RNA. The *gata2b* exon 4 CRISPR was synthesized as a crRNA and the ALT-R-CRISPR-Cas9 System (IDT) was used to make the injection mix. Briefly, crRNA was incubated with tracrRNA to form a gRNA complex followed by a second incubation with Cas9 protein (PNA Bio) to form the ribonucleoprotein complex (RNP). The RNP complex was then injected into embryos (50pg gRNA and 1μg protein). Founder fish screening and identification of F1 heterozygous adults by fluorescent PCR was performed as previously described^38^. All subsequent genotyping was performed using fluorescent PCR as described previously^36,38^. To amplify both the WT and mutant alleles in the same reaction for genotyping of *runx1* del(e3-8) mutants, we used a mixture of 4 primers as follows: a forward primer in exon 3, a forward primer in exon 8, common reverse primer in exon 8 and FAM-labeled M13F primer. This leads to the amplification of 280bp fragment in the WT fish (exon 8 primers) and 347bp fragment in the mutant fish (exon 3 forward and exon 8 reverse primers). To amplify both the WT and mutant alleles in the same reaction for genotyping of *gata2a* i4del194 mutants, we used a mixture of 4 primers as follows: a forward primer 5’ of the mutation, a forward primer within the deletion, common reverse primer 3’ to the mutation and FAM-labeled M13F primer. This leads to the amplification of 235bp and 452bp fragments in the WT fish and 255bp fragment in the mutant fish.

**Whole-mount *in situ* hybridization (WISH) and imaging**

WISH were carried out essentially as described by Thisse^39^. The following DIG-labeled antisense mRNA probes were generated by using UTP-digoxigenin (Roche): *c-myb*, *mpx*, *ae1-globin* (*hbae1*), *rag1*. Imaging and embryo observation was done using a Leica MZ16F stereo microscope equipped with a Leica DC7000T camera using Leica LFS 4.6.0. Images were collected using an Upright Zeiss Axio Imager D2 microscope (Carl Zeiss Inc., Thornwood, NY, USA).

**Histology and blood smears**

Blood smears were performed from 22-day-old larvae, euthanized with tricaine (MS-222), tail clipped and stained (Protocol Hema 3 system; Thermo Fisher Scientific). Paraffin sectioning (sagittal) and HE were performed by Histoserv, Inc. (Germantown, MD). Images were acquired using an AxioCam HRm CCD camera with a 1388 pixel x 1040 pixel imaging field. Zeiss ZEN blue pro 2011 software package was used for collection of all images.

**Kidney RNA extraction and RNA sequencing**

Wild type and *runx1^del25/del25^* kidneys were dissected from ~2.5 months old fish (n= 3 for each genotype, pool of 3 kidneys/replicate) and immediately collected in Trizol (#15596018, Invitrogen). Samples were then put in a heatblock at 52°C for 10 min and then overnight in -80°C, RNA was extracted following the manufacturer protocol using Direct-zol^TM^ RNA MiniPrep Kit (Zymo Research #R2051).

**Time lapse and confocal imaging**

Dechorionated embryos were anesthetized with tricaine, mounted in 0.8% low melting agarose and imaged using a Zeiss LSM 510 NLO Meta system(Thornwood, NY, USA) mounted on a Zeiss Axiovert 200M microscope with a Plan-Apochromat 20x/0.75 objective lens. Images were acquired every 10 minutes for a period of 15 hours. A range of 12-17 z-slices were used depending on the zebrafish orientation with a 1.94µm interval. Images were collected using the Zeiss ZEN 2009 V5.5 SP2 software package. Embryos were then recovered and genotyped according to the above protocol.

**FACS-sorting and single cell capture and sequencing with 10X Genomics Chromium**

Single cell suspensions were generated using a previously published protocol^44^ and sorted on a FACS AriaIII (Becton Dickinson, Franklin Lakes, NJ) using FACS DIVA software. Sorted cells were first gated on Forward Scatter Area versus Side Scatter Area dot plot. Two additional dot plots were used to remove cell clumps by using a singlet gate on Forward Scatter Area versus Forward Scatter Width followed by a second singlet gate on Side Scatter Area versus Side Scatter Width. Cells falling within these three gate regions were then evaluated for GFP^+^ cells on a dot plot of GFP (488nm laser excitation, emission 530/30nm bandpass) versus PE (488nm laser excitation, emission 576/26nm bandpass). GFP^low^ cells were collected in a small volume of DMEM-10%FBS (25-30 ul) in a PCR-tube and the final volume was then adjusted according to the 10X Chromium protocol.

**Single cell data processing, clustering and trajectory analysis**

For all our analysis raw data matrices of two or more Seurat objects were merged to generate a new Seurat object with the resulting combined raw.data matrix. The original identifiers for each dataset were set and only recovered at the end of the analysis. Cells with <200 and >4000 detected genes or with >5 percentage of UMIs mapped to mitochondrial genes were excluded from further analysis. The Seurat objects were processed according to the Seurat-guided clustering tutorial at <https://satijalab.org/seurat/vignettes.html>. Differential expression analysis between the wild type and mutant HSC/HSPC was done using Seurat default function “FindMarkers”. To perform single-cell trajectory reconstruction, we used STREAM software (Version 1.0)^17^ to analyze single-cell gene expression matrix exported from Seurat. In the dimension reduction step, we applied “se” method on “top_pcs” as features, and n_neighbors=50. In the elastic principal graph calculation step, parameters include epg_alpha=0.01, epg_mu=0.05, epg_lambda=0.01 were used. For other STREAM analysis procedurals, default parameters were used. Additional subway plot that reflect sample information and Seurat clustering information were generated with matplotlib (DOI: 10.1109/MCSE.2007.55). All R analyses were performed using R version 3.6.0. scRNA-seq are available at GEO under accession number #GSE158101.

**Mouse quantitative PCR**

Quantitative PCR (qPCR) was performed using Power SYBR Green PCR Master Mix (Applied Biosystems) according to manufacturer’s instruction. *Runx1* un-excised primers were used to detect un-excised Runx1 flox allele, and genomic control primers were used as internal controls for genomic DNA. The following primers were used: *Runx1* un-excised F 5’-ACTAGTGGATCCCTCGAGATAA-3’, *Runx1* un-excised R 5’-GGACTTGTTCTCCTGGTACAC-3’, Genomic control F 5’-CAGCCGTGGAGATTTAGGAG-3’, Genomic control R 5’-AGTGGGAGAACTGAGCGATG-3’, *Mx1-Cre* Genotype F 5’-CGATGCAACGAGTGATGAGG-3’, *Mx1-Cre* Genotype R 5’-GCATTGCTGTCACTTGGTCGT-3’, *Runx1* flox Genotype F 5’-CCCACTGTGTGCATTCCAGATTGG-3’, *Runx1* flox Genotype R 5’-GACGGTGATGGTCAGAGTGAAGC-3’.

**Statistical analysis**

Results are expressed as mean ± standard error of the mean. The statistical analysis was conducted using two-tailed Student’s t-test or a one-way ANOVA analysis using using GraphPad Prism v6.
