## Supplemental Figures 1-7 for "Redundant mechanisms driven independently by RUNX1 and GATA2 for hematopoietic development": Supplemental Figures.revised.pdf

### Supplemental figure 1

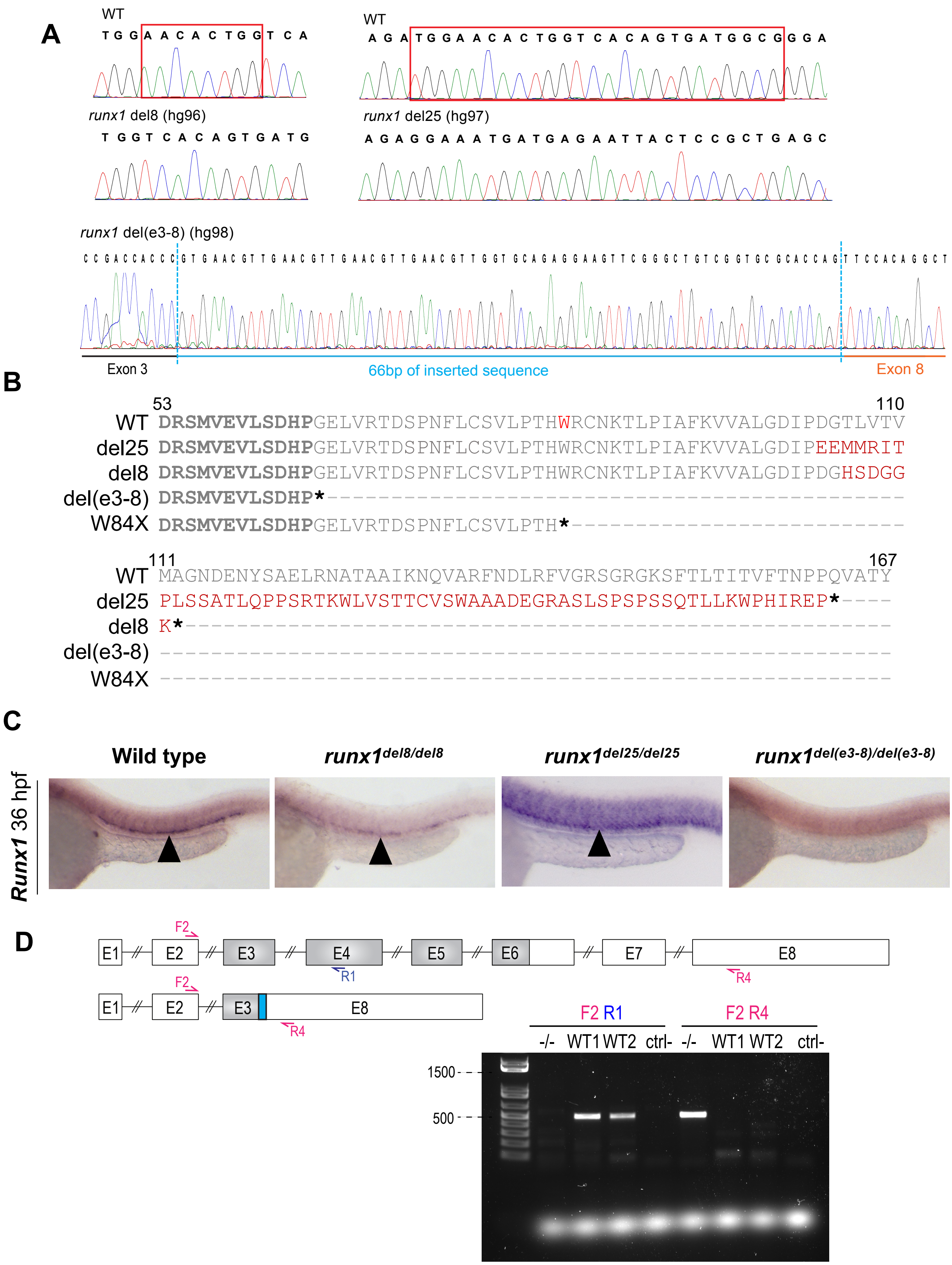

**Supplemental figure 1**

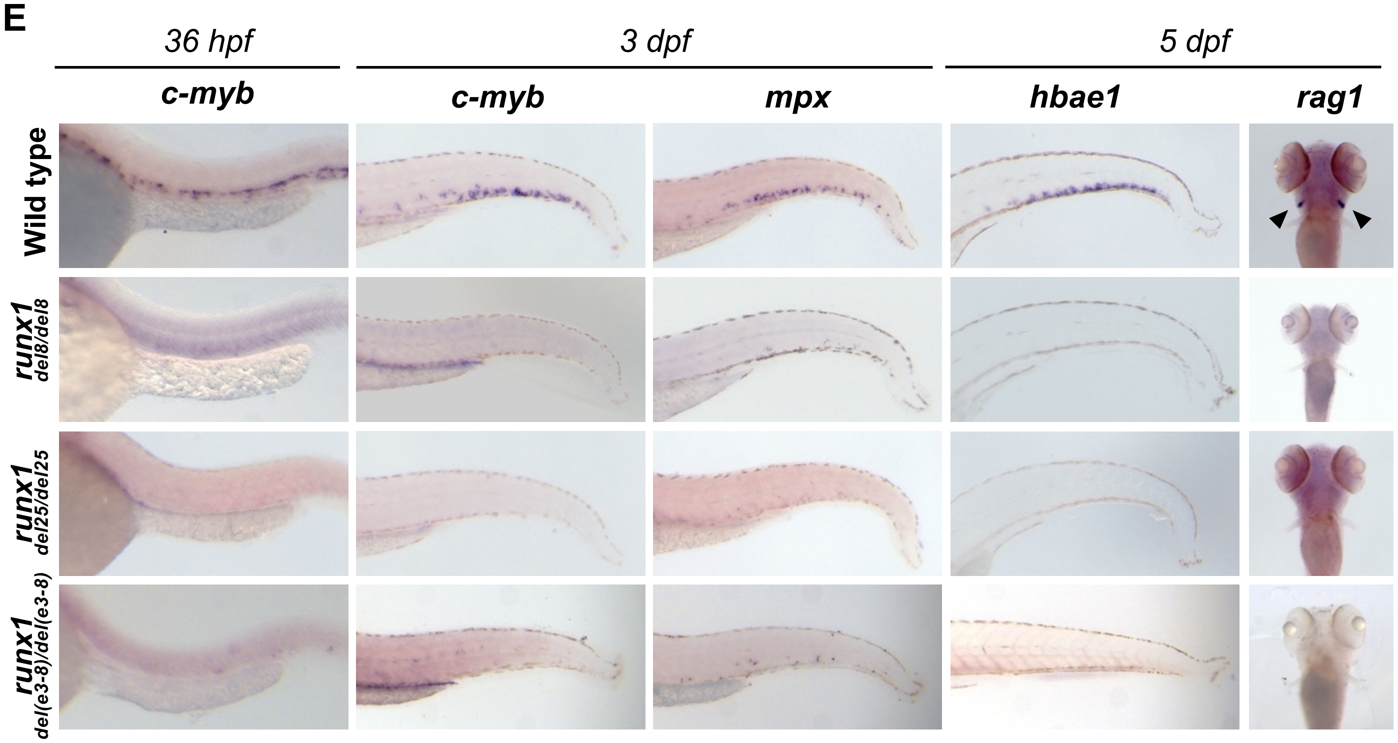


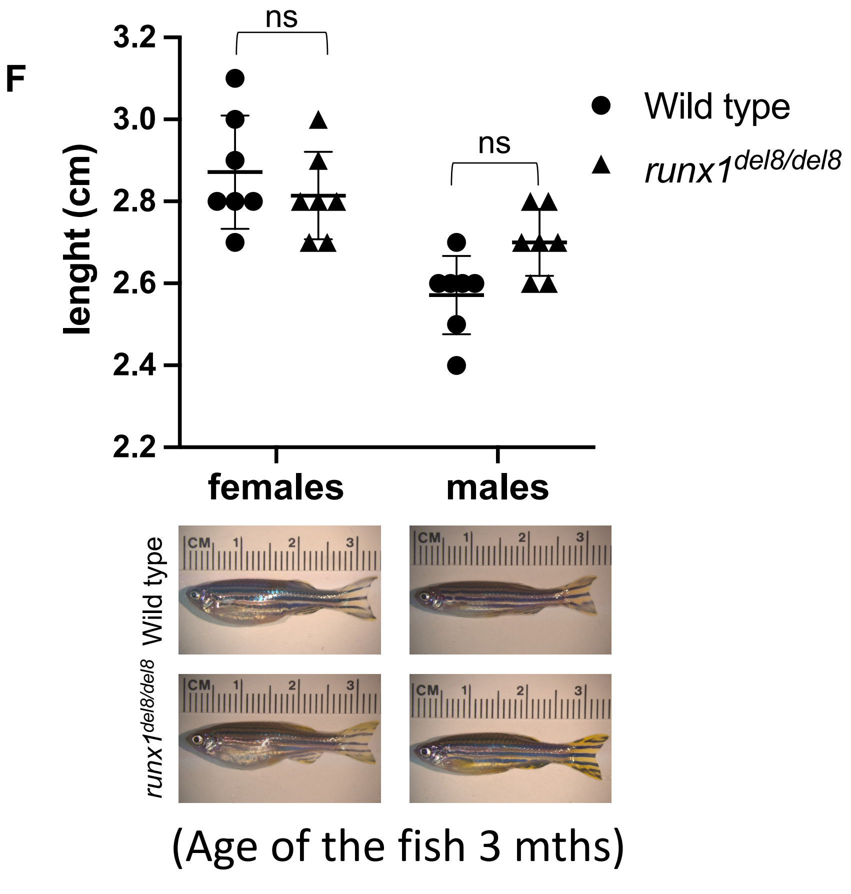


(Age of the fish 3 mths)

#### Supplemental figure 2

A

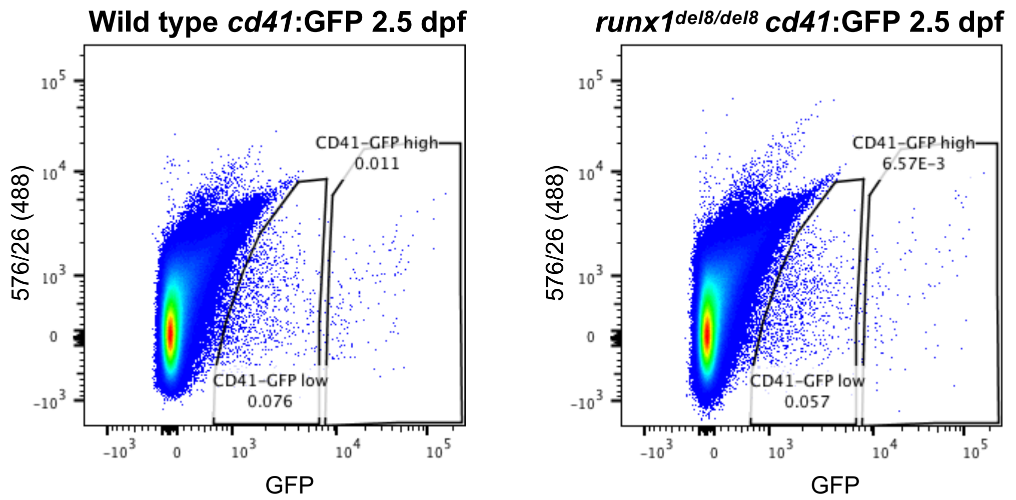


B

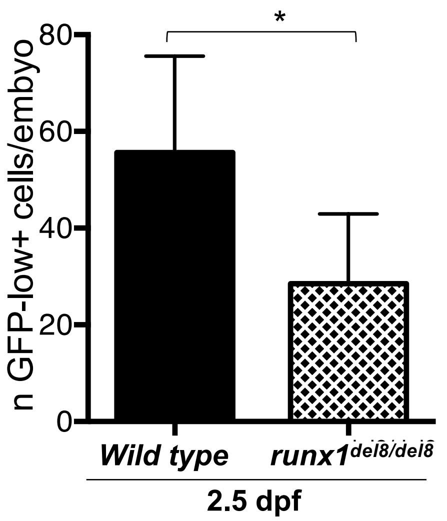


C

| *cd41*:GFP$^{low}$ | Wild type | *runx1*$^{del8/del8}$ |
|---|---|---|
| Total # of cells | 2200 | 1173 |
| Mean reads/cell | 112,873 | 251,483 |
| Median Genes/Cell | 1856 | 1258 |
| Median UMI/Cell | 12,290 | 7,757 |

D

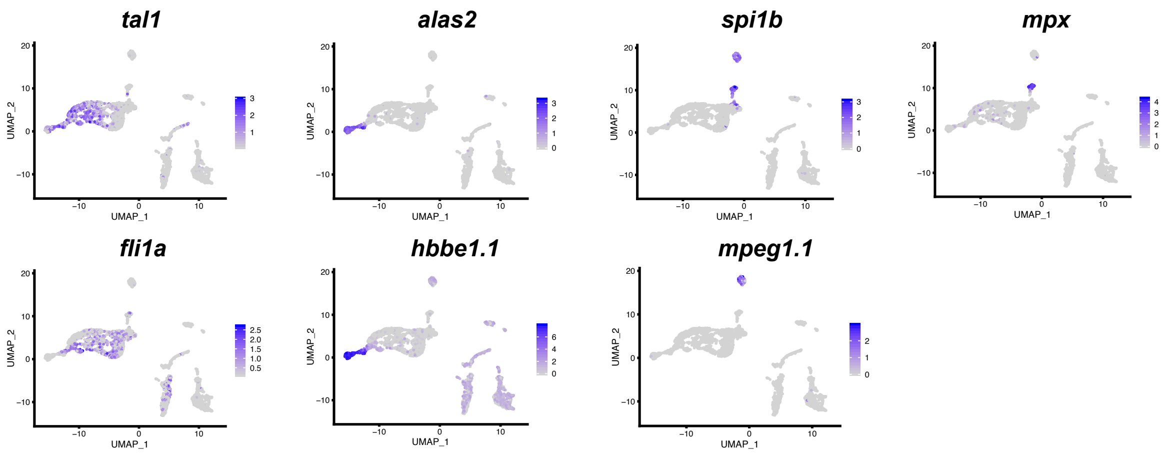

### Supplemental figure 3

**A**

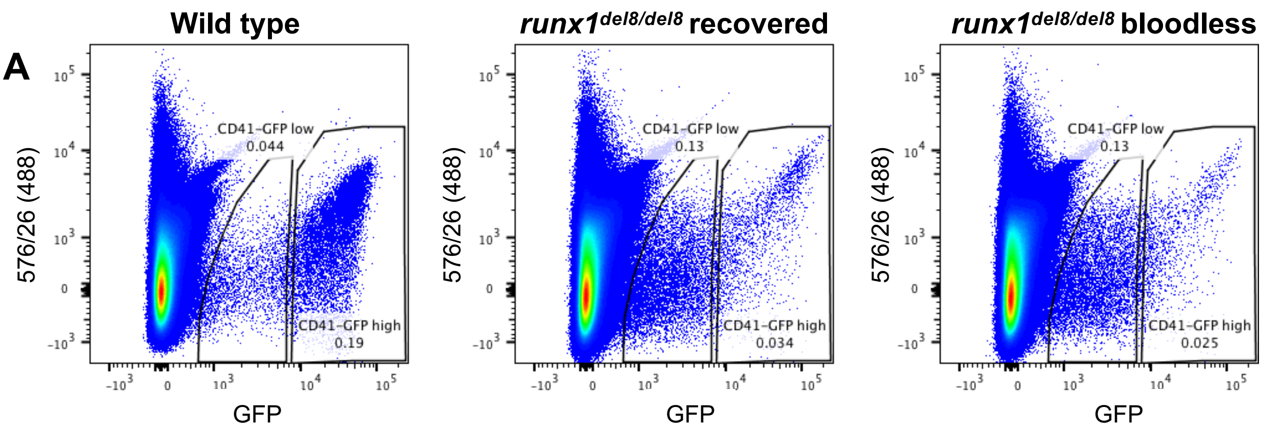


**B**

| *cd41*:GFP$^{low}$ | cd41-GFP$^{low}$ **6 dpf** | | cd41-GFP$^{low}$ **10 dpf** | | cd41-GFP$^{low}$ **16 dpf** | | |
|---|---|---|---|---|---|---|---|
| | ***Wild type*** | ***runx1$^{del8}$*** | ***Wild type*** | ***runx1$^{del8}$*** | ***Wild type*** | ***Runx1$^{del8}$ Bloodless*** | ***Runx1$^{del8}$ Recovered*** |
| **Total # of cells** | 2754 | 799 | 2711 | 3505 | 6218 | 3674 | 7883 |
| **Mean reads/cell** | 59341 | 226492 | 36914 | 33640 | 41619 | 64970 | 28464 |
| **Median Genes/Cell** | 1493 | 983 | 1731 | 1506 | 1769 | 1830 | 1713 |
| **Median UMI/Cell** | 13288 | 4054 | 10432 | 7145 | 12284 | 9372 | 10497 |

**C** 6 dpf

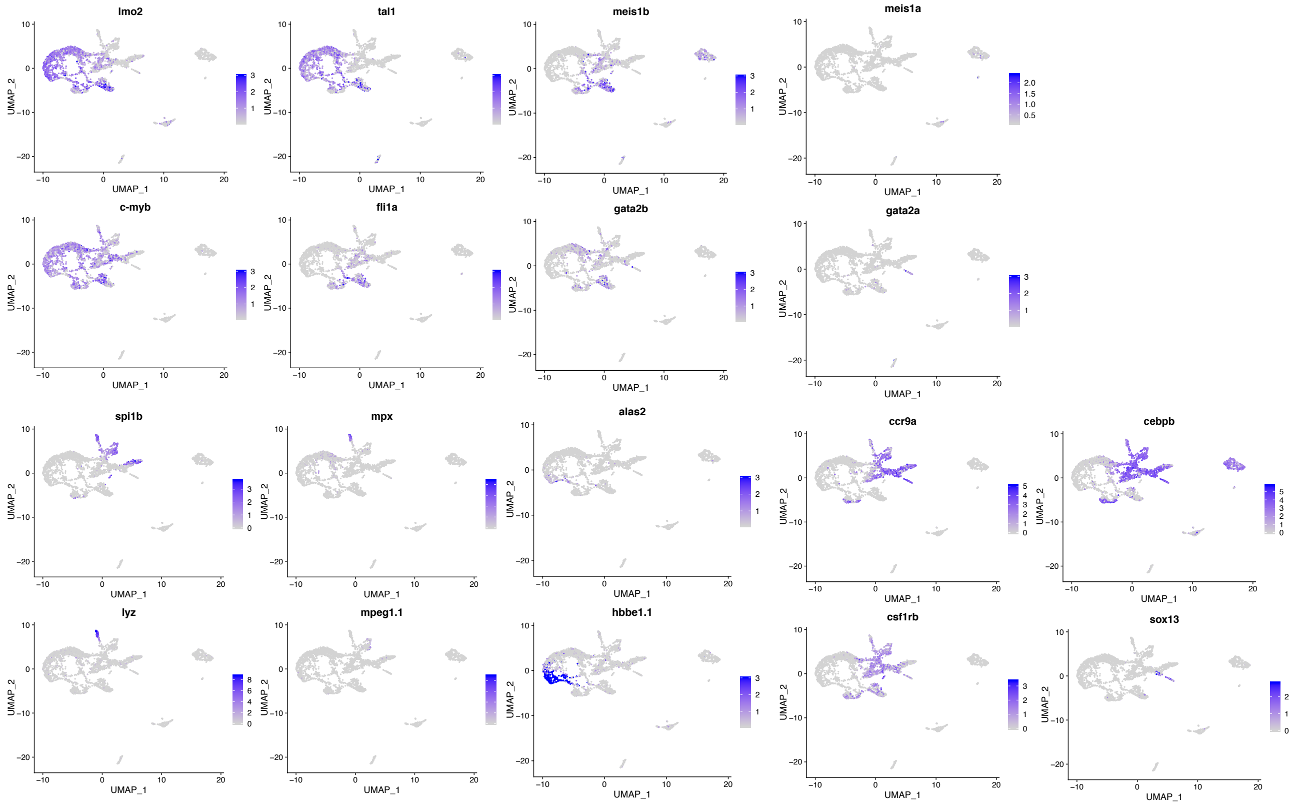


**D** 10 dpf

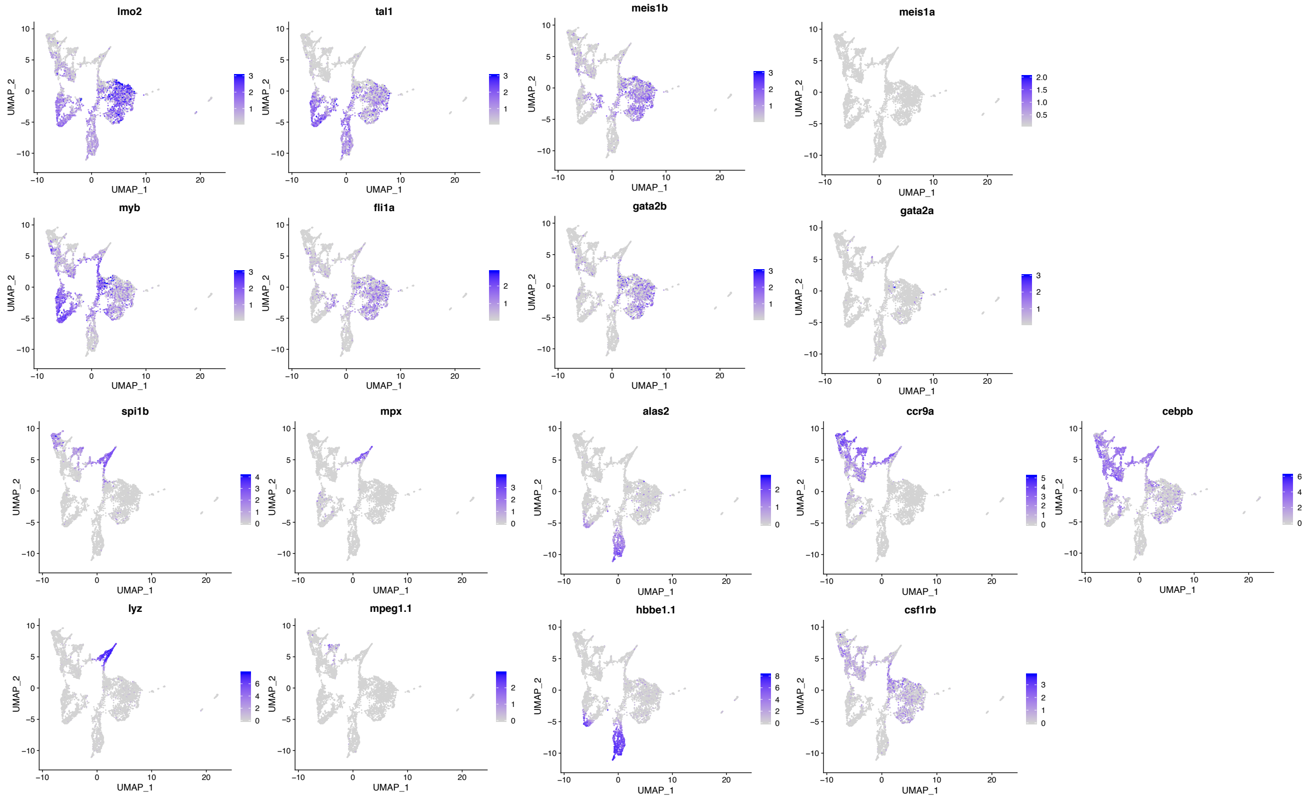

### Supplemental figure 4

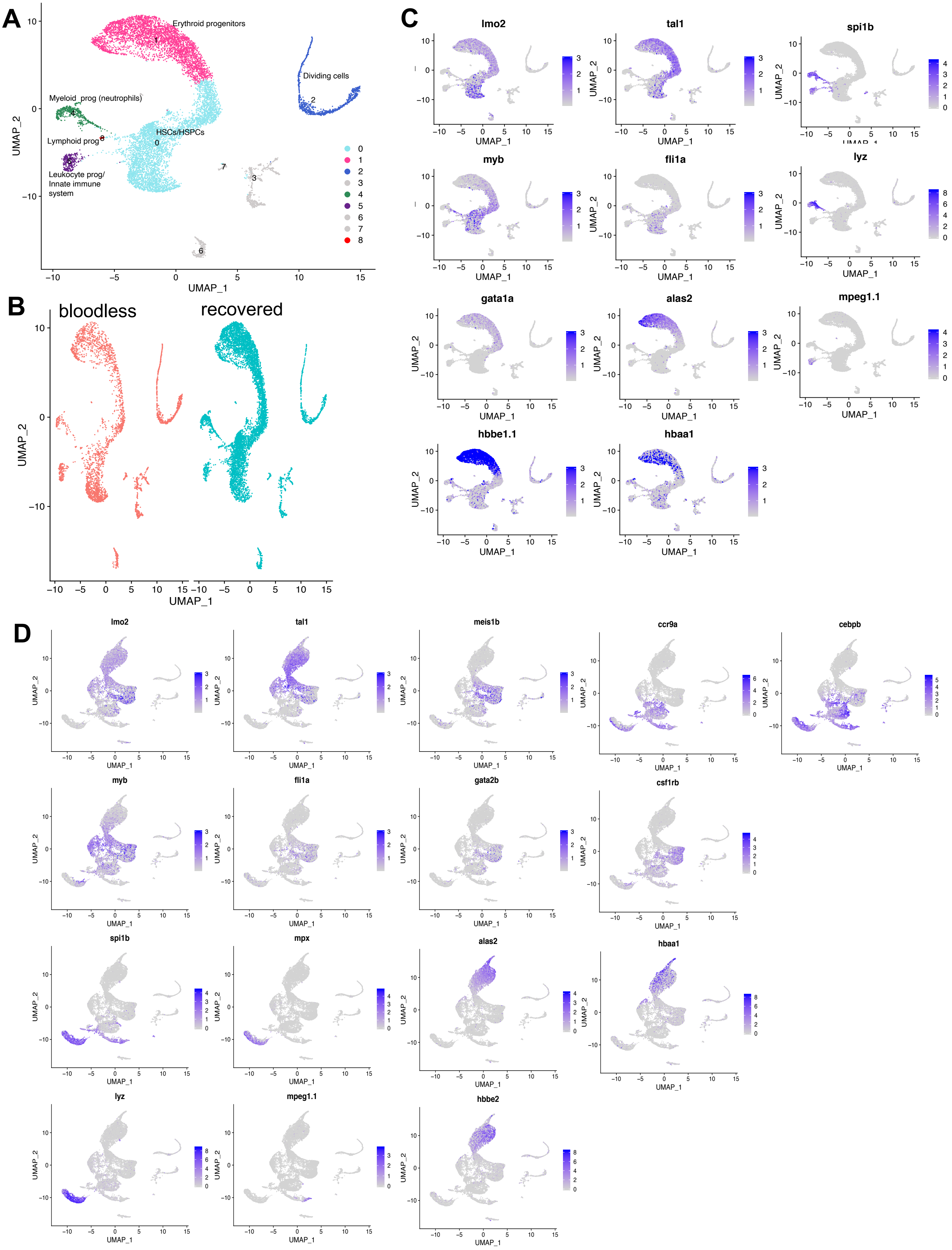

### Supplemental figure 5

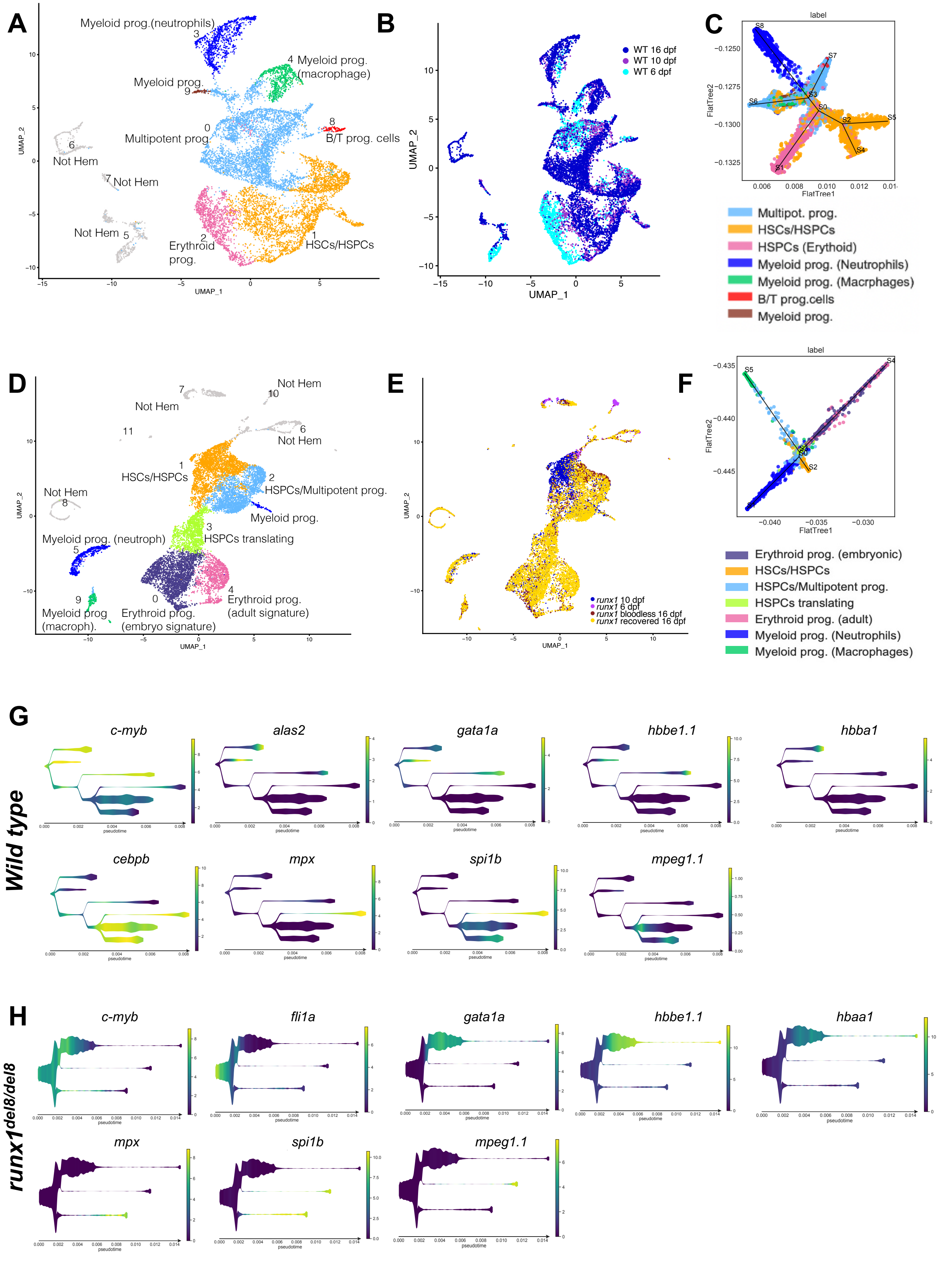

### Supplemental figure 6

A

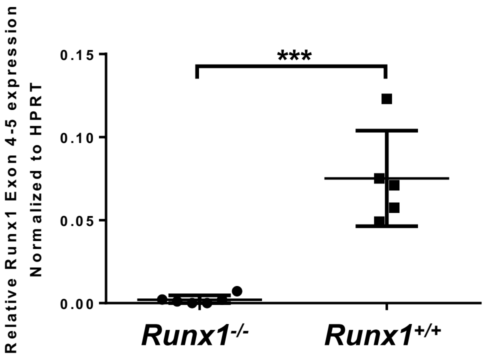


B

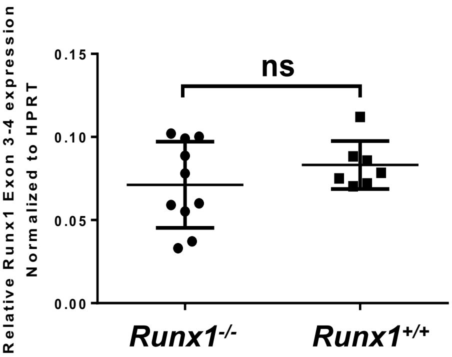


C

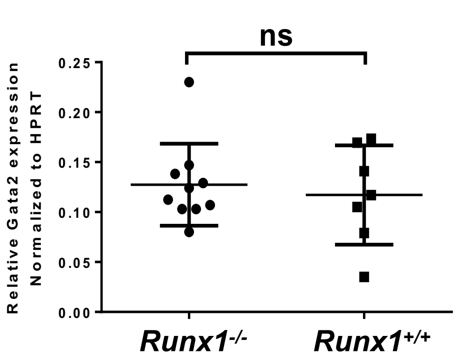


D

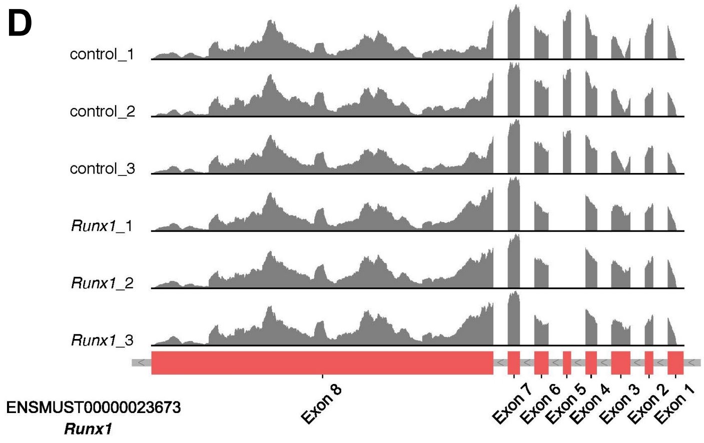


E

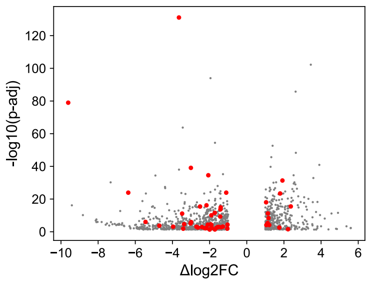


| DE genes c-Kit+ *Runx1* KO vs control | |
|---|---|
| Total | 1384 |
| UP | 414 |
| DOWN | 940 |

F

| *Gata2* targets | log2FC | padj |
|---|---|---|
| Mcpt8 | -3.650321322 | 8.02E-132 |
| Prss34 | -9.611277313 | 1.0477E-79 |
| Gpr141 | -3.008077068 | 7.9349E-40 |
| Fcnb | -2.069785301 | 2.6944E-35 |
| Gp5 | 1.919783526 | 4.4589E-32 |
| Fcer1a | -6.378849917 | 1.2049E-24 |
| Trem3 | -1.105839463 | 1.2049E-24 |
| Selp | 1.794588999 | 3.844E-24 |
| Runx1 | 1.04126559 | 9.5457E-19 |
| Ear2 | -2.168092334 | 6.7056E-17 |
| Treml1 | 2.369657329 | 3.193E-16 |
| Hba-a2 | -2.513858293 | 3.7829E-16 |
| Ms4a2 | -1.403797682 | 1.1998E-15 |
| E2f2 | -1.418846388 | 2.2369E-14 |
| Wfdc21 | -1.720175789 | 2.6872E-12 |
| Mfsd2b | 1.13729184 | 6.3402E-12 |
| I830127L07Rik | -3.478864935 | 7.8395E-12 |
| 4930438A08Rik | -1.907219749 | 8.3059E-11 |
| S100a8 | -1.428756293 | 5.2125E-10 |
| Il1rl1 | 1.170550668 | 4.6017E-09 |
| Il6 | -5.449056709 | 9.7035E-07 |
| Gm5900 | -2.99549301 | 1.1597E-06 |
| Gfi1b | 1.124317426 | 4.7116E-06 |
| Alas2 | -3.021702358 | 7.7742E-06 |
| Olfm4 | -3.37069312 | 2.4029E-05 |
| Serpinb10 | -1.939506569 | 2.7924E-05 |

| *Gata2* targets | log2FC | padj |
|---|---|---|
| Abca13 | -1.0365411 | 4.6065E-05 |
| Alox5 | -2.0995171 | 4.7741E-05 |
| Itgam | -1.905784 | 6.9314E-05 |
| F2rl2 | 1.2110646 | 7.4512E-05 |
| Aqp9 | 1.05463015 | 7.5652E-05 |
| Fcrla | -4.7062151 | 0.00018768 |
| Grap2 | -1.2854677 | 0.00068623 |
| Stfa1 | -3.9614682 | 0.00081374 |
| Cma1 | -2.7080113 | 0.00086956 |
| S100a9 | -1.5008933 | 0.00119023 |
| Cd96 | -1.550776 | 0.00119049 |
| Ngp | -2.409216 | 0.00138443 |
| Epx | -1.3739083 | 0.00163303 |
| Cd177 | -2.6266039 | 0.002584 |
| Tubb1 | 1.7535548 | 0.00287178 |
| Cd300ld | -1.8882208 | 0.00337986 |
| Mcemp1 | -1.6334332 | 0.00348528 |
| Camp | -2.1846944 | 0.00382526 |
| Lcn2 | -2.1893857 | 0.0052711 |
| Ltf | -2.2461519 | 0.00719916 |
| Trim10 | -1.0451479 | 0.01234823 |
| Gzmb | -3.4210466 | 0.01361247 |
| Mpig6b | 2.22068988 | 0.0295484 |
| Slc4a1 | -2.0049962 | 0.03829659 |
| Sytl3 | -1.9911064 | 0.04000622 |
| Mctp2 | -1.7169036 | 0.04123368 |

### Supplemental figure 7

**A**

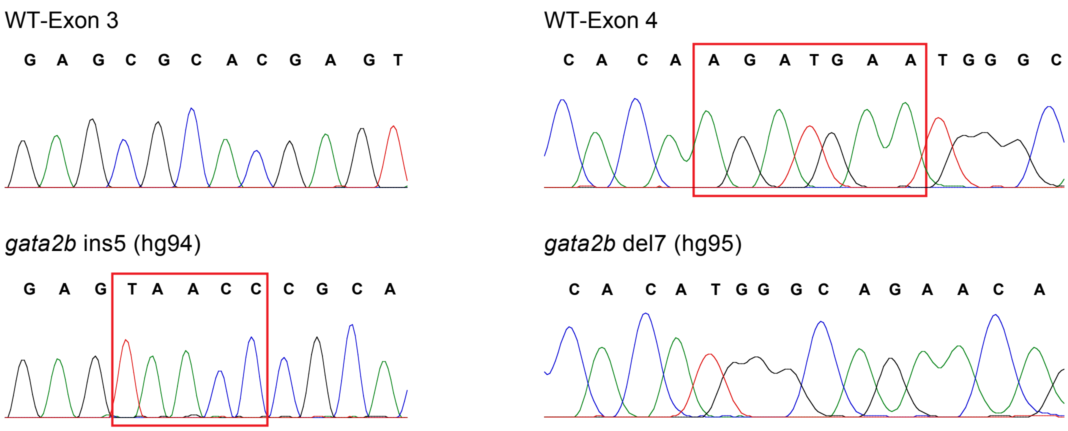


**B**

```
      220                                                         279
WT    PIPSYPDYSVAGAHEYPASVFHSRNLLGNMTTKCKSKNRAFSGRECVNCGATSTPLWRRD
ins5  PIPSYPDYSVAGVTRTSIPPVCSTPEICSET*----------------------------
del7  PIPSYPDYSVAGAHEYPASVFHSRNLLGNMTTKCKSKNRAFSGRECVNCGATSTPLWRRD

      280                                                         339
WT    GTGHYLCNACGLYHKMNGQNRPLIRPKRRLSASRRAGTCCANCQTGTTTLWRRNANGEPV
ins5  ------------------------------------------------------------
del7  GTGHYLCNACGLYHKGRTDLSSDPSADCQHLDEQAPVVPTARLGPPHSGDAMPTENPSAM

      340                                                         399
WT    CNACGLYYKLHNVNRPLTMKKDGIQTRNRKMSGKSKKRRGEHFHQFDSCVHDKPSSFSHM
ins5  ------------------------------------------------------------
del7  PAVYTTNYTM*-------------------------------------------------
```

**C**

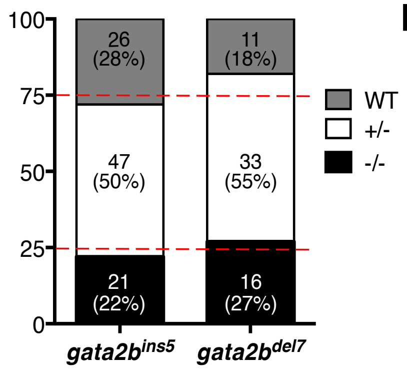


**D**

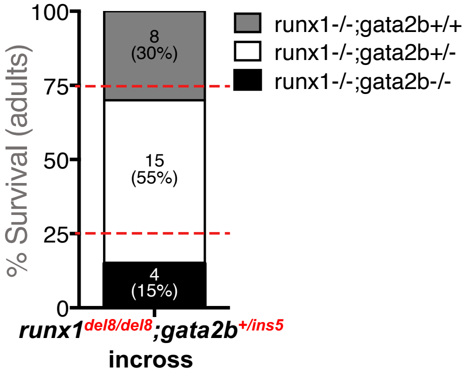

E

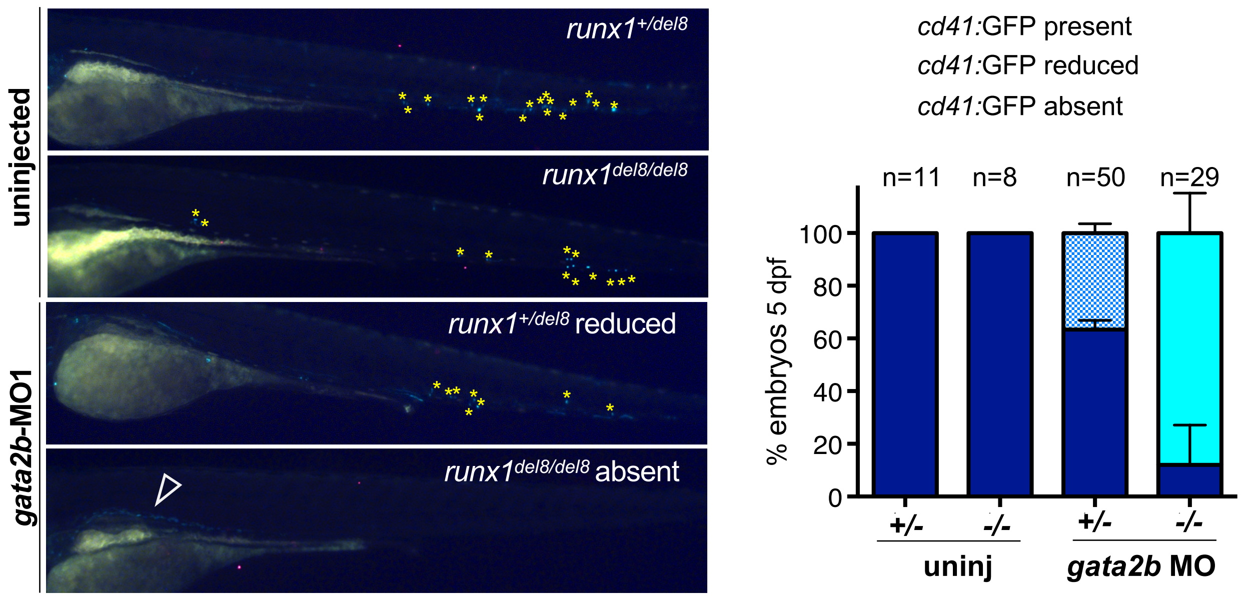
uninjected
gata2b-MO1
runx1+/del8
runx1del8/del8
runx1+/del8 reduced
runx1del8/del8 absent
cd41:GFP present
cd41:GFP reduced
cd41:GFP absent
n=11
n=8
n=50
n=29
% embryos 5 dpf
100
80
60
40
20
0
+/-
-/-
+/-
-/-
uninj
gata2b MO

F

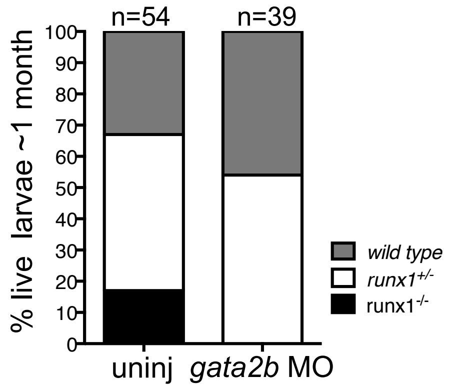
n=54
n=39
% live larvae ~1 month
100
90
80
70
60
50
40
30
20
10
0
uninj
gata2b MO
wild type
runx1+/-
runx1-/-

G

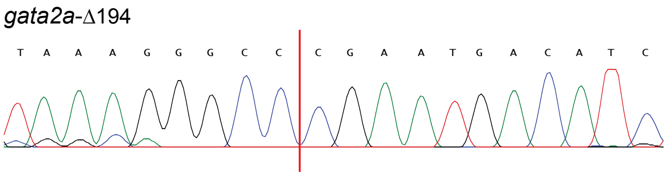
gata2a-Δ194
T A A A G G G C C C G A A T G A C A T C

H

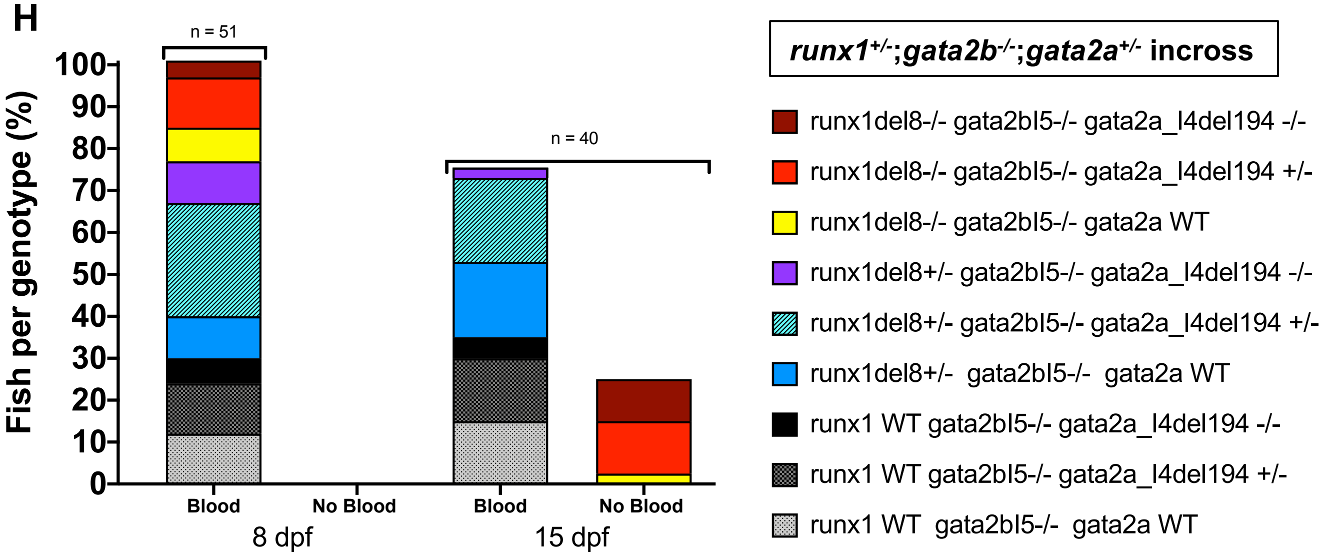
n = 51
n = 40
Fish per genotype (%)
100
90
80
70
60
50
40
30
20
10
0
Blood
No Blood
Blood
No Blood
8 dpf
15 dpf
runx1+/-;gata2b-/-;gata2a+/- incross
runx1del8-/- gata2bI5-/- gata2a_I4del194 -/-
runx1del8-/- gata2bI5-/- gata2a_I4del194 +/-
runx1del8-/- gata2bI5-/- gata2a WT
runx1del8+/- gata2bI5-/- gata2a_I4del194 -/-
runx1del8+/- gata2bI5-/- gata2a_I4del194 +/-
runx1del8+/- gata2bI5-/- gata2a WT
runx1 WT gata2bI5-/- gata2a_I4del194 -/-
runx1 WT gata2bI5-/- gata2a_I4del194 +/-
runx1 WT gata2bI5-/- gata2a WT
