## Supplemental Figures and Movies Legends for "Redundant mechanisms driven independently by RUNX1 and GATA2 for hematopoietic development"

**Supplemental Figure Legends and Movies Legends**

**Supplemental figure 1. Characterization of three new zebrafish *runx1* mutants.**

**A.** Sanger sequencing confirmation of the generated mutations. Upper panels show the presence of a deletion of 8 bp and a deletion of 25 bp in the *runx1* targeted with TALENs (*runx1^del8^* hg96 and *runx1^del25^* hg97). In the lower panel, the deletion in *runx1^del(e3-8)^* hg98 was confirmed, which was localized between exon 3 and 8, as well as a 66 bp insertion in the middle. **B.** Alignment of the WT RUNX1 protein sequence with the predicted protein sequences resulting from the three mutations: *runx1^del8^*, *runx1^del25^*, and *runx1^del(e3-8)^*. **C.** Expression of full length *runx1* mRNA can be detected in the AGM of *runx1^del8/del8^* and *runx1^del25/del25^* embryos at 36 hpf (black arrowheads) but not in the *runx1^del(e3-8)/del(e3-8)^* embryo by in situ hybridization. **D**. RT-PCR shows that the wild type *runx1* transcript is not detectable, but a mutant transcript is still present in the *runx1^del(e3-8)^* homozygous embryos (expected sizes of the PCR products: F2R1 (WT), 457 bp; F2R4 (del(e3-8)), 463bp). **E.** WISH showing the expression of definitive hematopoietic markers in the three *runx1* mutant lines and wildtype controls. The HSC marker *c-myb* is absent at 36 hpf in the AGM and at 3 dpf in the CHT. The myeloid progenitor marker *mpx* (3 dpf) and the erythroid marker *hbae1* (5 dpf) are also abrogated in the CHT of the *runx1* nulls. The lymphoid marker *rag1* is not detectable in the thymus (black arrowheads) of the *runx1^-/-^*. **F.** Size measurement of adult (3 months old) female and male wild type and *runx1^del8/del8^*.

**Supplemental figure 2. Data from single cell RNA-sequencing with 2.5 dpf embryos.**

**A.** Gating strategy for FACS isolation of *cd41*:GFP^low^ HSC/HSPC population from 2.5 dpf wild type and *runx1^del8/del8^* embryos. **B.** Counts of the *cd41*:GFP^low^ cells obtained from flow cytometric analysis of *Tg(cd41:GFP)* *runx1^del8/del8^* and wild type embryos at 2.5 dpf. For each experiment a pool of 25 - 130 embryos were analyzed for each genotype (n=4 wild type, n=6 *runx1^del8/del8^*). **C.**

Table listing single cell quality control metrics generated by 10X Cell Ranger. **D.** Feature plots showing the expression of selected markers (purple is high, grey is low) at 2.5 dpf. HSC: *tal1*, *fli1a*; erythroid EMPs: *alas2*, *hbbe1.1*; myeloid: *spi1b*; myeloid EMPs *mpeg1.1*, *mpx.*

**Supplemental figure 3. Data from single cell RNA-sequencing of *cd41*:GFP^low^ at larval stages .**

**A**. Gating strategy for FACS isolation of *cd41*:GFP^low^ HSC/HSPC population at larval stages. Data from 16 dpf wild type and *runx1^del8/del8^* embryos are presented and are representatives of all time-points. **B.** Table showing cell quality control metrics generated by 10X Cell Ranger. **C, D.** Feature plots showing the expression of selected markers (purple is high, grey is low) representing different populations at 6 dpf (C) and 10 dpf (D). HSC: *lmo2, tal1, c-myb, fli1a;* HSC mutant specific *meis1b, gata2b* are expressed but not *meis1a* or *gata2a.* Multipotent progenitors: *ccr9a, cebpb, csf1rb;* erythroid markers: *alas2, hbbe1.1.* Myeloid markers: *spi1b*, *mpx*, *mpeg1.1, lyz*; B/T prog cells: *sox13.*

**Supplemental figure 4. Single cell RNA-sequencing on 16 dpf larvae**

**A.** UMAP showing *cd41*:GFP^low^ cells from bloodless and recovered *runx1^del8/del8^* at 16 dpf. Colored clusters represent hematopoietic cells, grey clusters are non-hematopoietic (based on expression profile). **B.** UMAP showing the projection of the two genotypes merged for analysis. **C.** Feature plots showing the expression of selected markers (purple is high, grey is low) . HSC: *lmo2*, *tal1*, *c-myb*, *fli1a*; erythroid cells: *gata1a*, *alas2*, *hbbe1.1, hbaa1*; myeloid: *spi1b*, *lyz* and *mpeg1.1.* **D.** Feature plots showing the expression of selected markers (purple is high, grey is low) at 16 dpf (projection on UMAP presented in Fig. 3G). HSC: *lmo2, tal1, c-myb, fli1a;* expression of HSC mutant specific markers *meis1b, gata2b.* Multipotent progenitors: *ccr9a, cebpb, csf1rb;* erythroid markers: *alas2, hbbe1.1.* Myeloid markers: *spi1b*, *mpx*, *mpeg1.1, lyz.*

**Supplemental figure 5. Hematopoietic differentiation and pseudotime trajectories in *runx1^del8/del8^* and wildtype larvae at 6, 10, and 16 dpf**

**A.** UMAP showing merged wild type *cd41*:GFP^low^ cells from 6, 10 and 16 dpf. **B.** Distribution of the three time points used to generate the merged UMAP in figure A. **C.** Flat tree plot representing pseudotime trajectory projection and branches identities (STREAM) of wild type *cd41*:GFP^low^ cells from panel A. **D.** UMAP obtained by merging datasets from multiple larval timepoints 6, 10 and 16 (bloodless and recovered) dpf of *runx1^del8/del8^* *cd41*:GFP^low^  cells. In both A and D colored clusters represent hematopoietic cells, grey clusters are non-hematopoietic (based on expression profile). **E.** Projection of the timepoints on the UMAP in figure D. **F.** Flat tree plot showing the results of the trajectory analysis (STREAM) relative to UMAPs from *runx1^del8/del8^* (presented in panel D) merged dataset. **G.** Stream plots showing the expression of representative markers in wild types. HSC/HSPCs: *c-myb.* Erythroid markers: *alas2, gata1a, hbbe1.1*, *hbaa1*. Myeloid markers: *spi1b*, *mpx*, *mpeg1.1, lyz;* multipotent progenitors: *cebpb.* **H.** Stream plots showing the expression of representative markers in *runx1^del8/del8^*. HSC/HSPCs: *c-myb*, *fli1a.* Erythroid markers: *gata1a, hbbe1.1*, *hbaa1*. Myeloid markers: *spi1b*, *mpx*, *mpeg1.1.*

**Supplemental figure 6. GATA2 target genes are upregulated in c-Kit+ bone marrow cells from *Runx1* conditional knockout mice. A-C.** qPCR analysis on *Runx1^-/-^* (n=10) and *Runx1^+/+^* littermates (n=7) dissected AGM at E10.5. Expression of exon 5 is lost in *Runx1^-/-^* upon excision of the floxed allele (D) but the expression of the mutant mRNA appeared unchanged (E). Expression of *Gata2* in the *Runx1^-/-^* AGM was comparable to *Runx1^+/+^* (F). **D.** RNA-seq reads coverage plot of *Runx1* isoform ENSMUST00000023673 exon regions (three biological replicates) confirmed the complete excision of Exon 4 in all the *Runx1* knockout mice used for RNA-seq. **E, F.** The Volcano plot on the left displays the differentially expressed genes (padj <0.05, FC>2) between *Runx1* knockout group and control group, related to Figure 4B,C. Red dots represent the genes known to be regulated by the transcription factor GATA2 (H), which are also listed in the table on the right (G).

**Supplemental figure 7. Characterization of new zebrafish *gata2b* and *gata2a* mutants.**

**A.** Sanger sequencing shows the presence of an insertion of 5 bp (*gata2b^ins5^* hg94) in exon 3 and a deletion of 7 bp (*gata2b^del7^* hg95) in exon 4 of the *gata2b* gene. **B.** Predicted protein sequences of the mutant lines *gata2b^ins5^* and *gata2b^del7^*. **C.** Survival of zebrafish *gata2b^ins5^* and *gata2b^del7^* lines. Progenies of both lines were obtained from heterozygous incrosses.­ Red dashed lines indicate the expected Mendelian ratio. Segments on the bars show % of fish recovered for each genotype. The numbers in each segment depict the numbers of fish for each genotype. Adult *gata2b^-/-^* fish can be recovered according to Mendelian ratio. **D.** Survival of adult *runx1^del8^/gata2b^ins5^* double mutants obtained from the incross of *runx1^del8/del8^; gata2b^+/ins5^* parents. Red dashed lines indicate the expected ratio of *runx1^-/-^* recovery based on our previous experimental data (Fig. 1E). Each bar segment shows the percentage and number of fish recovered for each genotype. **E.** *gata2b* knock down experiments in the *runx1^del8^*; *Tg*(*cd41*:GFP) background. *cd41*:GFP^+^ cells were reduced in the *runx1^+/del8^* embryos and completely absent in the *runx1^del8/del8^* embryos after injection of *gata2b*-MO (9) (11.7 ng). Representative pictures of the phenotype at 5 dpf are shown in the left. Asterisks mark the *cd41*:GFP^+^ cells in the CHT, white arrowhead marks *cd41*:GFP^+^ cells in the pronephric duct. Right panel shows a quantification of the phenotype observed and the number of embryos analyzed. **F.** Survival of uninjected and *gata2b*-MO injected *runx1^del8/del8^* mutants (1 month old) obtained from incrossing *runx1^+/del8^* parents. 17% of the uninjected *runx1^del8/del8^* fish were recovered while no *runx1^del8/del8^* fish injected with *gata2b*-MO were recovered. **G.** Sanger sequencing confirming the deletion of 194 bp in the intron4 enhancer of *gata2a.* **H.** Presence or absence of blood circulation at 8 and 15 dpf in larvae obtained from *runx1^+/-^*;*gata2b^-/-^*;*gata2a^+/-^* incross. Segments on the bar show % of fish recovered for each genotype. Bloodless larvae are observed only in presence of *runx1^-/-^*.

**Supplemental Movie 1.** ***cd41:*GFP^+^;*kdrl:mCherry^+^* cells emerge and are released from the AGM through the axial vein in wild type embryo *Tg(cd41:GFP); Tg(kdrl:mCherry)*.**

Time lapse imaging of the AGM region of a wild type embryo at 2.5 dpf. mCherry marks endothelial cells, GFP is expressed in thrombocytes, HSCs and non-hematopoietic cells. HSCs derived from hemogenic endothelium are mCherry^+^GFP^+^ (yellow).

Images were acquired every 10 minutes for a period of 15 hours. A range of 12-17 z-slices were used depending on the zebrafish orientation with a 1.94µm interval.

**Supplemental Movie 2. *cd41:*GFP^+^;*kdrl:mCherry^+^* are present in the *runx1^-/-^* AGM and are released through the axial vein in a *Tg(cd41:GFP); Tg(kdrl:mCherry) embryo*.**

Time lapse imaging of the AGM region of a *runx1^-/-^* embryo at 2.5 dpf. mCherry marks endothelial cells, GFP is expressed in thrombocytes, HSCs and non-hematopoietic cells. HSCs derived from hemogenic endothelium are mCherry^+^GFP^+^ (yellow).

Images were acquired every 10 minutes for a period of 15 hours. A range of 12-17 z-slices were used depending on the zebrafish orientation with a 1.94µm interval.
